# Foraminiferal environmental DNA reveals late Holocene sea-level changes

**DOI:** 10.1101/2025.08.12.669806

**Authors:** Zhaojia Liu, Nicole S. Khan, Howard K.Y. Yu, Arthur Chung, Magali Schweizer, Celia Schunter

## Abstract

Reconstructing past relative sea level (RSL) provides critical insight into the mechanisms driving RSL change and informs future projections. Foraminifera are widely used sea-level indicators, but their application is often limited by poor preservation. Here, we demonstrate that foraminiferal environmental DNA (eDNA) and sedimentary ancient DNA (sedaDNA) provide a complementary approach to traditional morphological methods for RSL reconstruction. By analyzing surface sediments and a core from subtropical intertidal environments in the Pearl River Delta, we found that foraminiferal eDNA and sedaDNA assemblages exhibit clear vertical zonation, consistent with morphological results. An eDNA-based transfer function enabled high-resolution RSL reconstructions with decadal temporal and decimeter vertical precision from two periods: 290–1703 CE and 1956–present. Notably, eDNA preservation extended RSL reconstruction far earlier than morphological analyses (1956 CE). The eDNA reconstruction closely matched tide-gauge and geological RSL records, underscoring its potential as a robust tool for reconstructing past RSL and its driving mechanisms.

## 1.1 Introduction

Holocene relative sea level (RSL) records extend knowledge of mechanisms driving RSL change beyond the limited duration instrumental record^1–4^. These records enable comparison with other reconstructed climatic variables and model-based predictions to identify drivers of RSL change and their interactions within the climate system^5–8^ and thus inform future sea-level projections^4,9^.

Holocene RSL reconstructions are derived from geological proxies that exhibit systematic and quantifiable relationships to tidal elevation and thus RSL^10^. Among these, salt-marsh foraminifera are widely used due to their (1) ubiquity in low-energy intertidal environments across the globe; and (2) low-diversity assemblages found in high numbers^11,12^. Their sensitivity to subaerial exposure (inundation frequency) results in distinct vertical zonation within the tidal frame^13–15^. Knowledge of their present-day distribution and ecological preferences can be used to interpret assemblages of fossil foraminifera preserved in sediment archives to reconstruct past tidal elevation and produce high-resolution (centimeter to decimeter-scale) records of RSL change^16–18^. However, a major limitation of this approach is the poor or selective preservation of foraminiferal tests, particularly in (sub)tropical settings where elevated temperatures and precipitation promote test degradation or dissolution^19–21^. This problem in part arises from the traditional morphological methods used to identify and enumerate foraminiferal assemblages, which rely on the identification of individuals under a stereomicroscope based on test morphology.

Environmental DNA (eDNA), DNA obtained from environmental samples (soil, sediment, air, and water) without isolating or culturing target organisms, offers a promising alternative^22^. eDNA metabarcoding using a fragment of the small subunit of ribosomal RNA gene (SSU rDNA) is increasingly being used in foraminifera studies, including environmental biomonitoring^23–26^ and biodiversity assessments^24,25,27^. Compared to traditional morphological methods, eDNA reveals greater diversity in modern benthic foraminifera, especially for organic-walled or finely-agglutinated monothalamids^28^. After foraminifera die, their DNA binds to environmental compounds, such as clay minerals, humic acids and other charged particles, providing protection from degradation by microbial DNases (deoxyribonucleases) in the environment^29–32^. Additionally, the relatively anoxic and radiation-free environment in buried sediments facilitates the preservation of this sedimentary ancient DNA (sedaDNA) for hundreds to thousands of years^33–35^. As such, sedaDNA has been used in foraminiferal paleoecological and paleoceanographic reconstructions^36–41^. However, these reconstructions have primarily focused on open-ocean or fjord environments with cold and/or anoxic conditions that differ from the low-energy intertidal settings that produce precise records of RSL. Despite its potential, foraminiferal eDNA remains unexplored as a sea-level proxy.

Here we evaluate the utility of foraminiferal eDNA and sedaDNA as a sea-level proxy in intertidal environments from subtropical Hong Kong. We characterize contemporary foraminifera vertical distribution using eDNA metabarcoding and morphological methods from three transects and examine the environmental factors controlling their distribution. To better reconcile taxonomy between eDNA and morphological methods, we develop an in-house DNA sequence database of dominant taxa identified through morphological analysis. We use these contemporary assemblages to interpret a radiocarbon-dated mangrove core. A Bayesian transfer function (BTF) is developed from the modern eDNA training set, incorporating prior information from the elevation range of environmental zones identified by stable carbon isotope geochemistry, to reconstruct paleo-mangrove elevation and RSL change. Despite limitations with the preservation quality of sedaDNA in sediment layers lacking foraminiferal tests, this approach provides robust reconstructions of paleo-mangrove elevation (PME) and RSL change that are consistent with regional tide gauge and RSL datasets. These findings establish foraminiferal eDNA and sedaDNA as a valuable tool for paleoenvironmental and sea-level studies.

### 1.2 Study site

Samples were collected from a site where foraminifera, diatoms, and geochemical proxies and chronological analyses were previously examined for paleoenvironmental and RSL reconstructions^42,43^ (Fig. 1). The site was chosen due to its proximity to a tide gauge and availability of historical aerial imagery showing mangrove and mudflat extent, enabling validation of reconstructions. Previous investigations identified foraminifera through morphological analyses conducted across three transects and sediment core MPSC01^43^. Six environmental zones— subtidal, mudflat, mangrove fringe, mangrove, terrestrial transition, and terrestrial— were differentiated based on δ^13^C, total organic carbon (TOC), and carbon to nitrogen (C/N) ratios geochemistry (Supplementary Fig. 1; Supplementary Table 1). Together with radiocarbon dating of the core, these proxies revealed a transition from tidal flat organic mud to mangrove peaty mud and muddy peat at 1957 CE (Supplementary Table 2).

**Fig. 1.**
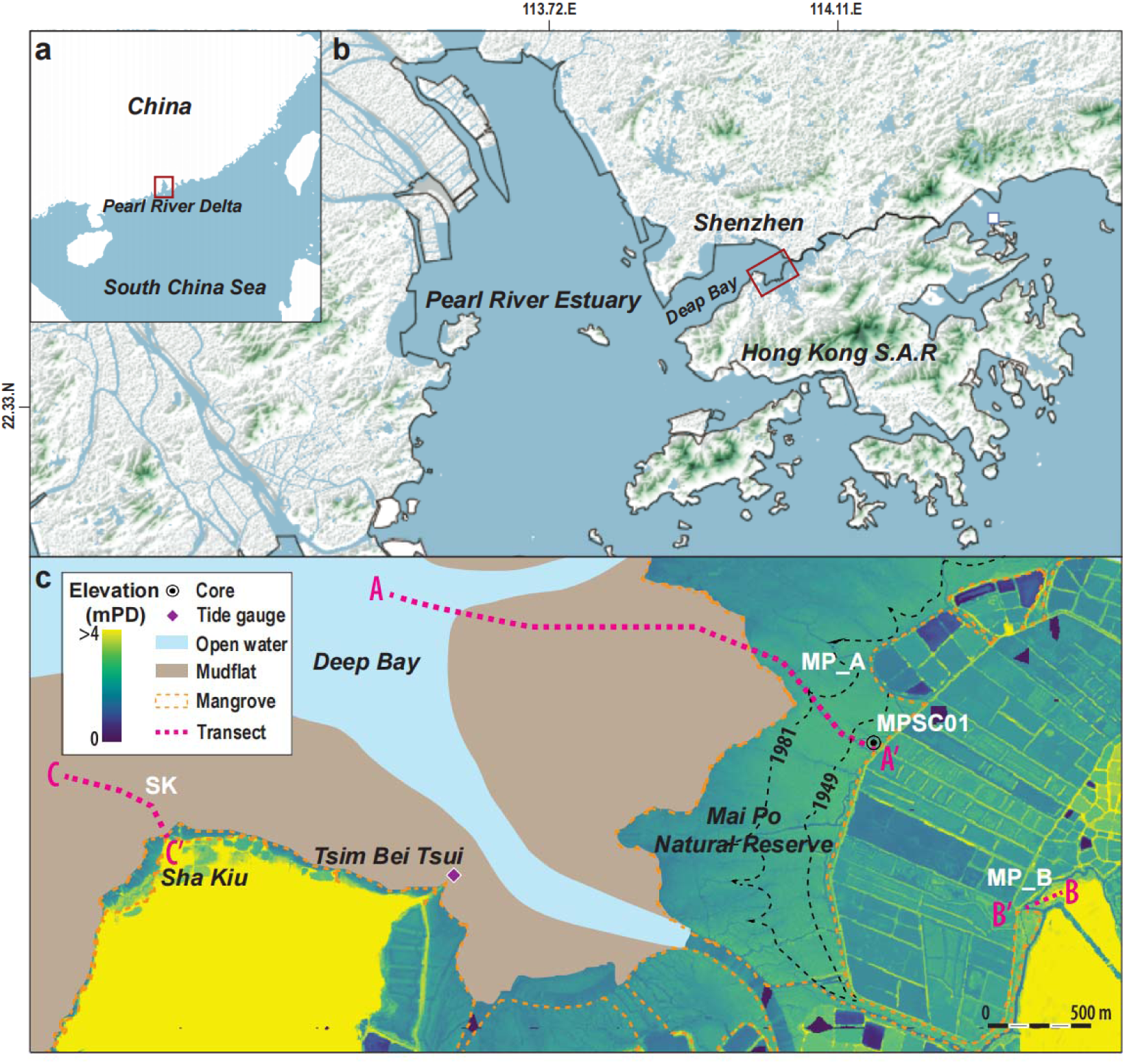
Location of the study area in the Pearl River Delta (PRD) (a) and in Deep Bay (b). Location of surface sampling transects (MP_A, MP_B, and SK), sediment core MPSC01, the nearest tide gauge at Tsim Bei Tsui, and the land surface elevation as represented by the LiDAR digital elevation model provided by the Hong Kong S.A.R government in Mai Po and Sha Kiu (c). Historical boundaries of mangrove in Mai Po Nature Reserve inferred from aerial images^43^ are indicated by the black dashed line.

## 2. Results

### 2.1 Foraminiferal eDNA assemblages

The foraminiferal eDNA/sedaDNA dataset revealed substantial genetic diversity, encompassing 824 operational taxonomic units (OTUs) derived from 20,897,581 reads from 60 modern and 28 core samples after filtering out non-foraminiferal, low-quality, and chimeric reads. Taxonomic resolution varied, with 22 OTUs identified at the species level (9% of total sequences), 4 at the genus level (0.1%), 327 at the family level (48%), and 4 at the order level (0.2%) based on GenBank comparisons. An additional 9 OTUs were assigned to species level using an in-house database developed by sequencing abundant local taxa absent in public databases (Supplementary Table 3). The remaining 458 OTUs were identified as foraminifera (Phylum Foraminifera), but could not be assigned to higher taxonomic levels (Supplementary Table 4).

Of the assigned OTUs, 165 belonged to calcareous taxa, 98 to agglutinated taxa, and 95 to monothalamids. The most abundant hard-shelled families were Ammoniidae and Miliamminidae, collectively representing 33% of the total sequences, while other hard-shelled families (e.g. Trochamminidae, Elphidiidae) contributed fewer (7%) sequences (Supplementary Fig. 2, 3; Supplementary Table 4). Monothalamous OTUs (e.g., Allogromiidae and Saccamminidae) contributed 19% of total sequences. Undetermined foraminiferal OTUs accounted for 37%, with two dominant OTUs (OTU3 and OTU6) contributing 12% (Supplementary Fig. 2, 3; Supplementary Table 4).

Distinct assemblages were observed in modern samples across the elevation gradient, expressed in standardized water level index units (SWLI) that normalize tidal elevations across sites with different tidal characteristics. At high elevations above 100 SWLI (equivalent to mean tide level), Miliamminidae dominated (relative abundances of sequences ranging from 1-93%) along with undetermined OTUs (e.g. OTU3: ≤43%; OTU11: ≤17%), Ammoniidae (≤42%), and Trochamminidae (≤14%) (Fig. 2). Ammoniidae were prevalent (1-55%) in upper mudflat and mangrove environments (57–200 SWLI), while Nummulitidae (1-23%) and OTU6 (≤ 73%) dominated low-elevation mudflat and subtidal environments (<57 SWLI). Monothalamous families exhibited broad elevation tolerance, occurring across subtidal to mangrove environments (Allogromiidae: ≤15%; Saccamminidae: ≤62%) (Fig. 2c).

**Fig. 2.**
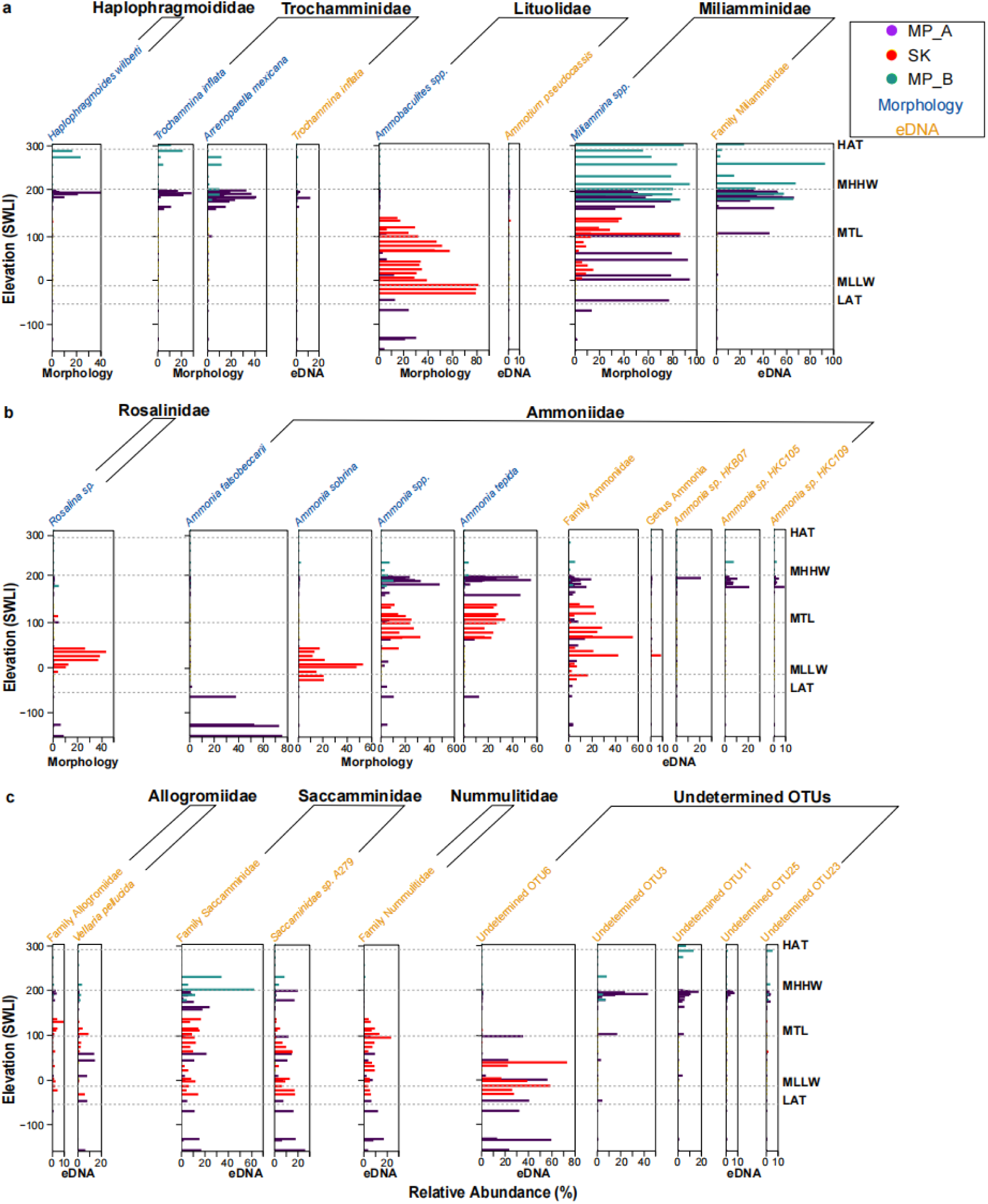
Vertical distribution of major taxa in the morphological and eDNA assemblages, separated by agglutinated (a), calcareous taxa (b) and taxa in eDNA assemblage without counterparts in the morphological assemblage (c). Elevation is presented as Standardized Water Level Index (SWLI) units, of which a value of 100 represents mean tide level (MTL) and a value of 200 represents mean higher high water (MHHW).

### 2.2 Morphological foraminiferal assemblages

A total of 8,660 individuals representing 24 morphospecies were identified from modern and core samples, including 14 agglutinated and 10 calcareous taxa across 12 families (Supplementary Fig. 2, 3). Seven families from the morphological dataset linked to 194 OTUs that contribute 35% of the total reads (33% in modern samples and 44% in core samples). Five morphospecies from five different families—*Acostata* sp., *Bruneica* sp., and *Glomospirella* spp. (lacking reference sequences in GenBank), as well as *Haplophragmoides wilberti* and *Rosalina* sp.—were not detected in the eDNA/sedaDNA assemblage (Supplementary Fig. 2, 3).

Similar to eDNA/sedaDNA, Miliamminidae and Ammoniidae were dominant, with Miliamminidae present in 92% of modern samples (Fig. 2a), contributing 41% and 53% of total abundance in modern and core samples, respectively (Supplementary Fig. 2, 3). However, Miliamminidae sequences were rare below MTL. Ammoniidae exhibited similar distribution patterns in mangrove environments across both datasets (Fig. 2b). Some taxa abundant in the morphological assemblages (e.g., *Ammonia sobrina*, absent from GenBank, and *A. tepida* and *A. falsobeccarii*) were not found in eDNA data, causing discrepancies in low-elevation environments (Fig. 2b).

Low-elevation subtidal and mudflat environments (<57 SWLI) were dominated by calcareous taxa, including Ammoniidae (e.g., *A. falsobeccarii*: ≤76% in each sample; *A. sobrina*: ≤53%), Rosalinidae (≤43%), and agglutinated Lituolidae (≤81%), collectively accounting for 68% of counts in these environments (Fig. 2a, 2b). In upper mudflat and mangrove fringe environments (60–136 SWLI), Lituolidae (≤56%), Ammoniidae (*Ammonia* spp.; 5–32%; *A. tepida*: ≤ 34%) dominated, together contributing 71% of total counts (Fig. 2a, 2b). At higher elevations (>136 SWLI), agglutinated Trochamminidae (*A. mexicana*: ≤ 41%; *T. inflata*: ≤ 29%) and Haplophragmoididae (≤40%) became increasingly abundant (Fig. 2a).

### 2.3 Environmental variables shaping morphological and eDNA assemblages

Canonical correspondence analysis (CCA) and partial CCA (pCCA) revealed salinity, δ^13^C, and elevation as key environmental factors shaping eDNA assemblages, explaining 20%, 16%, and 15% of the variance, respectively (Fig. 3a, 3b; Supplementary Tables 5, 6). In comparison, elevation (23%), followed by sand content (17%) and salinity (13%) explained the variability in morphological assemblages (Supplementary Table 6). Non-metric multidimensional scaling (NMDS) using Bray-Curtis dissimilarities of community composition revealed elevation preferences of taxa from both morphological and eDNA assemblages (Fig. 3c, 3d). The community composition was significantly influenced by elevation, with strong effects observed for both morphological (PERMANOVA; F = 33.43, R² = 0.366, *p* = 0.001) and eDNA assemblages (F = 27.49, R² = 0.325, *p* = 0.001).

**Fig. 3.**
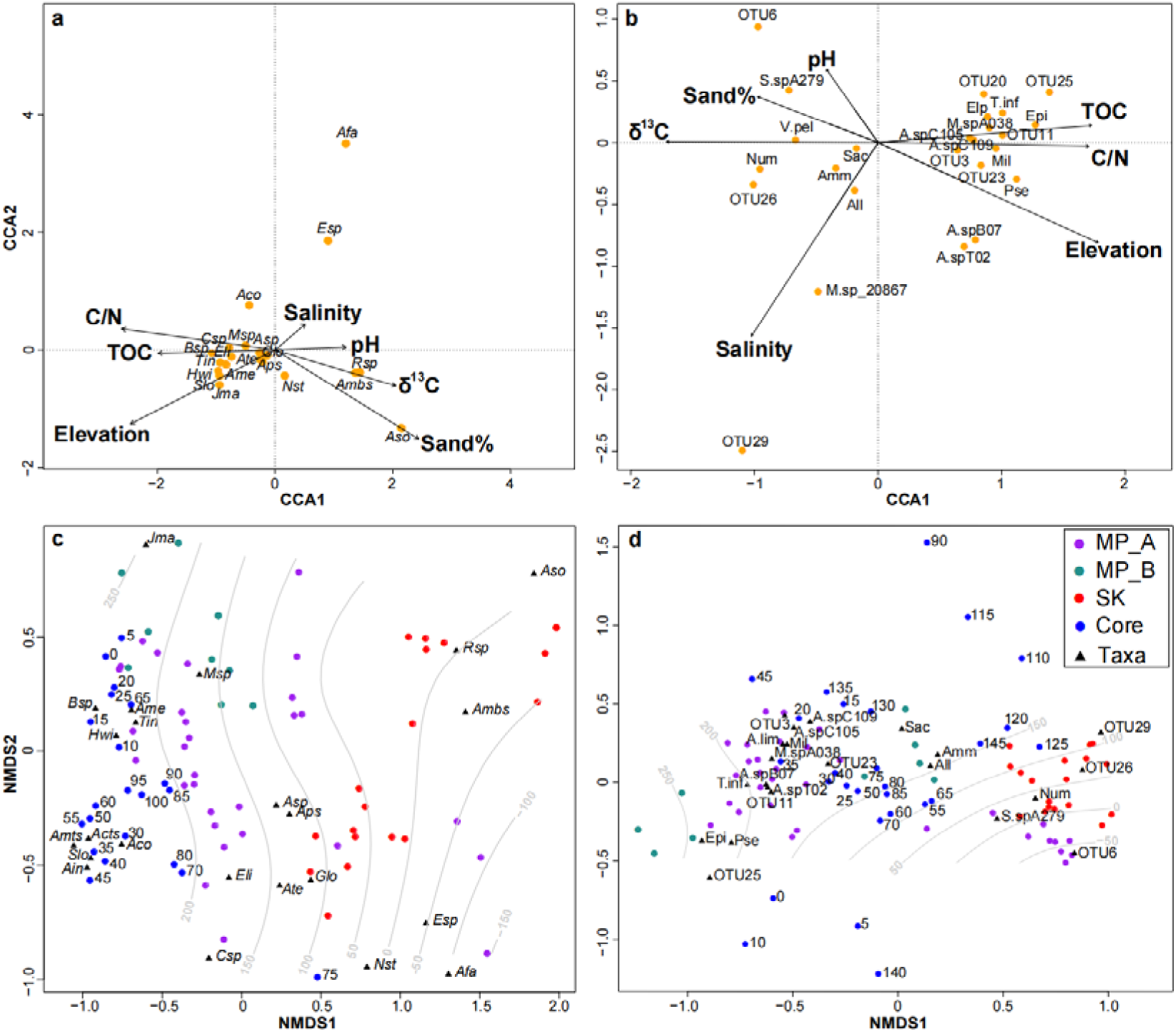
Multivariate ordination analyses conducted on morphological and DNA assemblages reveal the influence of examined environmental variables on taxa composition. Taxa-environment Canonical Correspondence Analysis (CCA) biplots of morphological (a) and DNA (b) taxa that exceed 5% of the assemblage in at least one sample. The lengths of the environmental arrows on the taxa-environment biplot approximate their relative importance of influence on the foraminiferal assemblage. Arrow direction reflects its approximate correlation with the ordination axes and other environmental factors, and the perpendicular projection of taxa position on the environmental arrow indicate their weighted average optima along each environmental variable. The first two axes of eDNA and morphological CCA explained 16% and 37% of total variance, respectively. Non-metric Multi-Dimensional Scaling (NMDS) ordination of morphological (c) and DNA (d) assemblages of all modern surface sampling stations and core samples based on Bray-Curtis dissimilarities. Samples collected from MP_A, MP_B and SK transect were marked with different colors. Core samples are shown as depth below surface (cm). Grey lines indicate the relative standard water level index (SWLI) elevations associated with sample positions in NMDS ordination space. Some major undetermined OTUs and assigned taxa of eDNA assemblage are shown. Taxa names are written in abbreviation (Supplementary Tables 7, 8).

### 2.4. Downcore signatures of foraminiferal eDNA and morphological assemblages

Three lithostratigraphic units (Unit I-III) were identified in core MPSC01 (Fig. 4). Foraminiferal sedaDNA was successfully extracted and amplified from sediments above 0.85 m PD, indicating sufficient preservation to permit paleoenvironmental interpretation. Nevertheless, the observed OTUs in the sedaDNA assemblage was significantly lower than those in the modern eDNA assemblage (ANOVA; *p* <0.05). Units I and II displayed similar α-diversity metrics, as measured by observed OTUs, Shannon, and Chao1 indices (Supplementary Table 9). In contrast, the sedaDNA assemblage in Unit III exhibited significantly lower observed OTUs and Chao1 indices (ANOVA; *p* <0.05), suggesting the potential influence of taphonomic processes with depth.

**Fig. 4.**
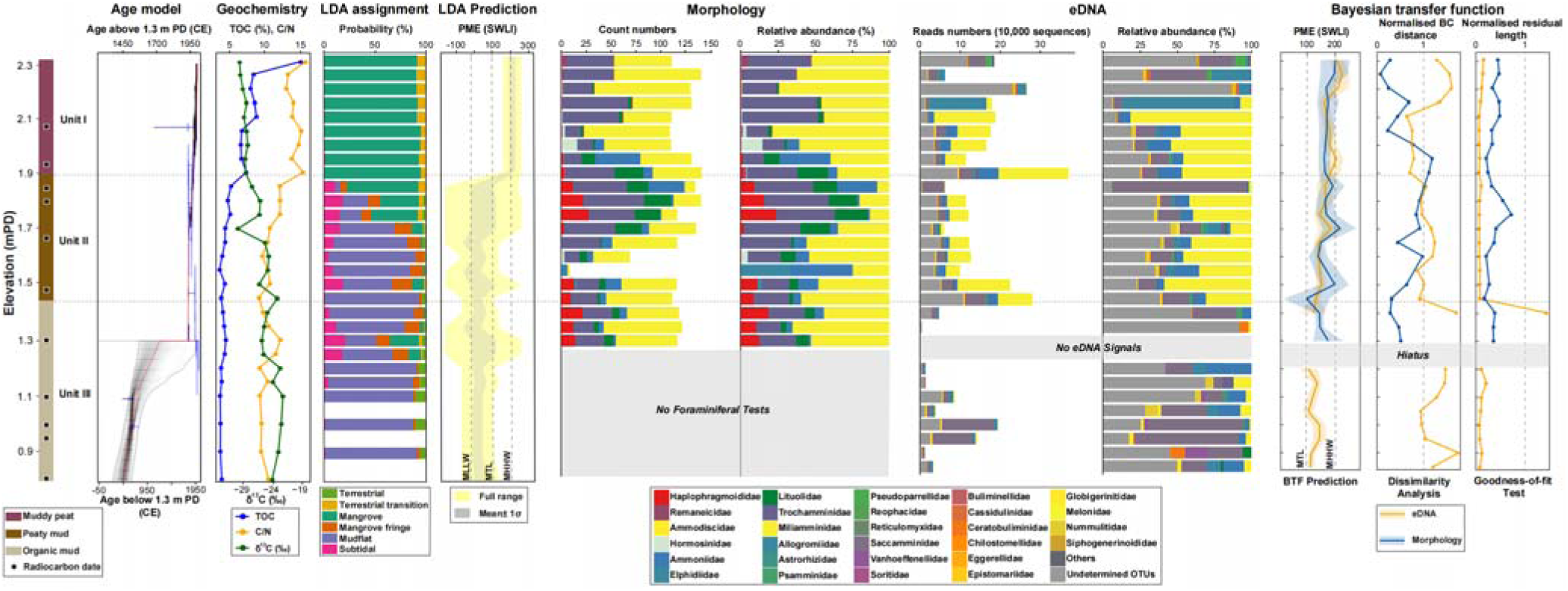
Core MPSC01 lithology, chronology established by Bacon age-depth model (age outliers were removed but shown in Supplementary Fig. 4), geochemistry variables and associated linear discriminant analysis (LDA) prediction of environmental zones, paleo-mangrove elevation (PME) prediction inferred from LDA, foraminiferal morphological and sedaDNA assemblages, reconstructed PME estimated by foraminiferal eDNA- and morphology-Bayesian transfer function (BTF), dissimilarity index based on Bray-Curtis (BC) distance, and goodness-of-fit statistics. Morpho- and eDNA taxa are grouped at the family level for comparability. “Others” includes OTUs assigned to order level. The envelop of BTF predictions represent ±20-uncertainties. For the dissimilarity index, the 10th percentile of the sedaDNA datasets and the 20th percentile of the morphological datasets was normalized to 1, with values >1 suggesting poor analogy to a modern counterpart. For goodness-of-fit statistics in core samples, the 95% threshold of squared residual lengths was normalized to 1. Values below 1 indicate a poor fit of the transfer function to elevation.

Foraminiferal sedaDNA reveals distinct changes in core assemblages with depth. The sedaDNA assemblage in the basal lithostratigraphic unit (Unit III; 0.85–1.44 m PD)—deposited between 486 (290-652, 2σ uncertainty range) CE and 1957 (1957-1958) CE—resembled the modern mudflat eDNA assemblage, characterized by low Miliamminidae abundance (≤17%), moderate Ammoniidae (≤30%), and substantial Saccamminidae (6-69%) contributions (Fig. 4). These patterns align with linear discriminant analysis (LDA) applied to core geochemistry, which also indicated a mudflat environment (Fig. 4).

In Unit II (1.45 to 1.89 m PD), which was deposited between 1957 and 1991 CE, sedaDNA assemblages were dominated by Miliamminidae (14-82%), with moderate contributions from Ammoniidae (2-16%) and Saccamminidae (2-18%) (Fig. 4). This shift, characterized by increased Miliamminidae sequences, reflects a transition from a low-elevation mudflat to a mangrove environment and is consistent with LDA predictions (Fig. 4). An exception at 1.85–1.86 m PD showed Saccamminidae dominance (91%), which lacked modern analogues and likely indicates a localized disturbance or depositional change at this boundary between lithostratigraphic units.

Similarly, the sedaDNA assemblages in Unit I (1.89 to 2.15 m PD), deposited after 1991 CE, were consistently dominated by Miliamminidae (46–81%), a pattern characteristic of modern mangrove environments and supported by LDA results (Fig. 4). However, in samples near the top of the core (above 2.15 m PD), Miliamminidae abundance dropped (<8%), while Saccamminidae (≤38%) and undetermined OTUs (e.g., OTU25 ≤58%) increased. This assemblage lacks a modern analogue in the training set and may reflect influence by bioturbation.

Compared to the sedaDNA assemblage, the morphological assemblage preserved in the core exhibited a consistent composition across all lithostratigraphic units (1.30–2.31 m PD), being dominated by agglutinated taxa, including Miliamminidae (8–76%) and Trochamminidae (*A. mexicana*, 10 – 39%; *T. inflata* ≤ 20%) (Fig. 4). This composition reflects a mangrove environment, aligning with interpretations from the sedaDNA assemblage. However, although no significant difference in α-diversity metrics was observed between modern and core morphological assemblages (ANOVA; *p* >0.05), taphonomic issues in the core were more evident than with the sedaDNA assemblage. Foraminiferal tests were entirely absent below 1.30 m PD in Unit III (deposited before 1956 CE), and tests were sparse (<100 individuals per 1 cm³ sediment) between 1.55 and 1.61 m PD (Fig. 4), preventing paleoenvironmental interpretations of sedimentary archive based solely on morphological assemblages.

### 2.5 Performance of eDNA and morphological training sets

We developed Bayesian transfer functions (BTFs) from eDNA and morphological training sets to estimate response curves (mean with 95% credible interval) for each taxon in relation to tidal elevation (SWLI). Many taxa in the eDNA assemblage exhibited unimodal distributions with tidal elevation, including several abundant OTUs (e.g. OTU26 and OTU29) and monothalamous species (e.g. *Vellaria pellucida*). However, some taxa in both assemblages displayed bimodal or multimodal distributions, including abundant families Saccamminidae and Ammoniidae (Fig. 5b, 5c).

**Fig. 5.**
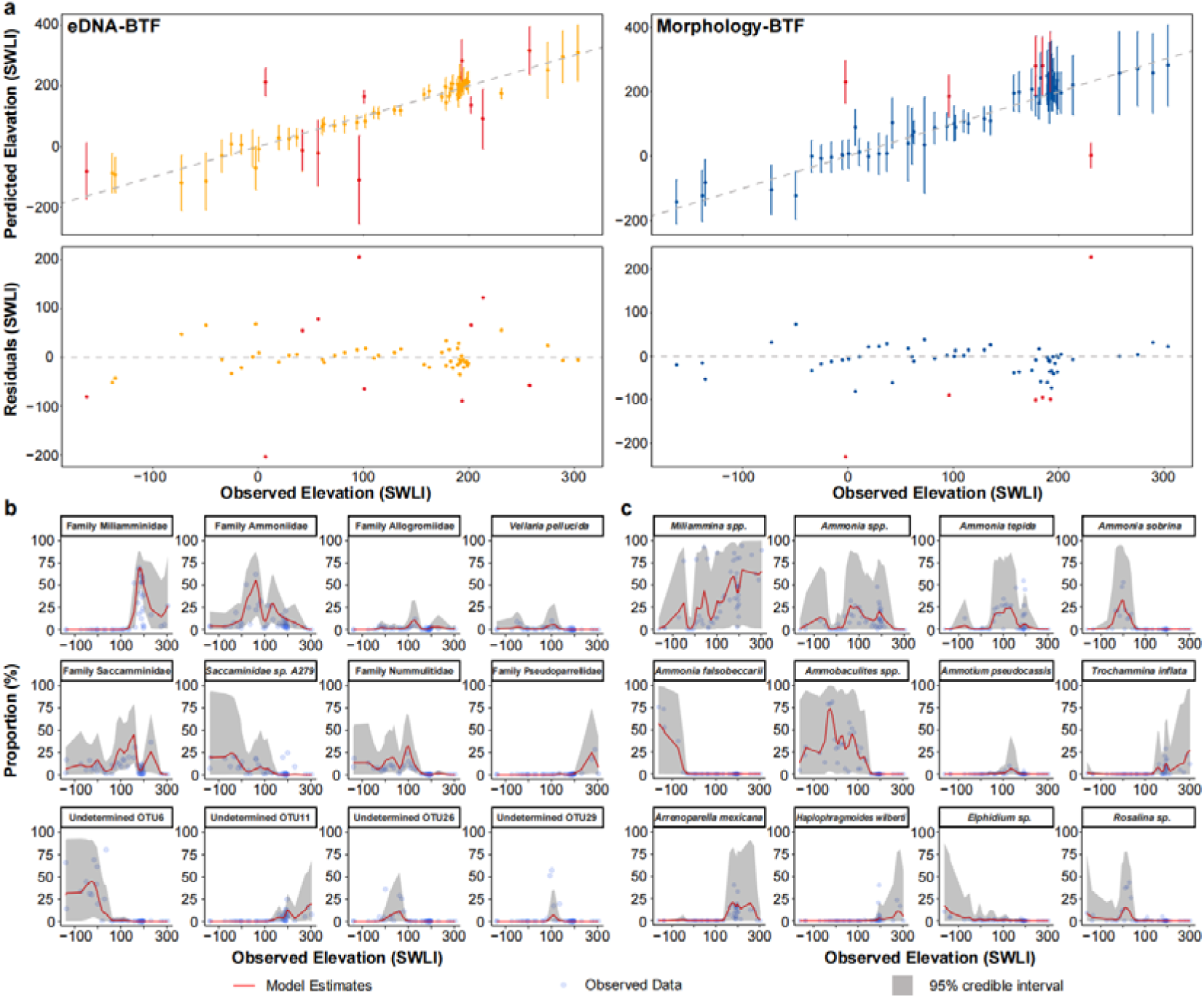
Ten-fold cross validation and taxa response curves modeled by the Bayesian transfer function (BTF). BTFs predicted elevation by 10-fold cross-validation versus observed elevation measured at sampling stations of foraminiferal eDNA and morphological assemblages modern training set are shown in (a). Elevation was expressed as standardized water level index units (SWLI). Residuals were calculated by subtracting predicted value from observed elevation. For both assemblages’ BTFs, samples that were screened out due to residuals exceeding two standard deviations from the mean residual in cross-validation are also shown, but highlighted in red. Taxa response curves of major taxa from eDNA (b) and morphological (c) assemblages. Circles represent the relative abundance of corresponding taxa at observed sample elevations, the red solid line and the grey shaded envelope represent the model mean and the 95% credible interval of prediction.

To evaluate BTF performance, a 10-fold cross-validation was conducted on both eDNA and morphological datasets (Fig. 5; Table 1; Supplementary Table 10). Both BTFs showed strong correlations between observed and predicted elevation (R² = 0.95 for eDNA, 0.93 for morphology). After removing outliers, the morphology-BTF slightly outperformed the eDNA-BTF in accuracy (98% vs. 94%), but the eDNA-BTF had lower uncertainty (1σ = 18 vs. 34 SWLI) and RMSEP (25 vs. 32). Elevated uncertainty and outliers were mainly found in the low and high ends of the SWLI range for the eDNA-BTF, and the morphology-BTF tended to overpredict elevation in lower-mudflat environments (Fig. 5a).

**Table 1.** Summary of eDNA- and morphology-Bayesian transfer functions (BTFs) performance in 10-fold cross-validation of the modern training set.

| Metric | eDNA-BTF | Morphology-BTF |
| --- | --- | --- |
| Correlation ($R^2$) | 0.95 | 0.93 |
| $1\sigma$ Uncertainty (SWLI) | 18 | 34 |
| RMSEP | 25 | 32 |
| Accuracy (within $2\sigma$ uncertainty range) | 94% | 98% |
| Average Residual (SWLI) | 0.7 | -9 |

### 2.6 Reconstructed paleoenvironmental and paleo-mangrove elevation change

The eDNA-BTF predicted an increase in paleo-mangrove elevation (PME) from 131 to 210 SWLI from core base to top, with an average 1σ uncertainty of 12 SWLI (equivalent of 0.5 m). In comparison, the morphology-BTF estimated a rise in PME from 179 to 204 SWLI from 1.3 m PD to core top, with an average 1σ uncertainty of 18 SWLI (0.6 m) (Fig. 4).

To evaluate the reliability of our reconstruction, we compared it with annual mean sea level (MSL) changes in Hong Kong recorded by tide gauges since 1963 CE and other sea-level data from the Pearl River Delta (PRD) (Fig. 6). Excluding core samples exceeding dissimilarity index thresholds, the eDNA-BTF estimated a RSL rise of −0.01 to 0.08 meters between 1978 and 2016 CE (Fig. 6a), capturing all tide gauge records within 2σ uncertainties. This trend was consistent with the morphology-BTF, but with a slightly higher mean squared error (MSE: eDNA = 0.027 m²; morphology-BTF = 0.016 m²) (Fig. 6c). The eDNA-BTF extended the RSL record back to 486 (290-652) CE, supported by index and limiting points from the PRD (Fig. 6d).

**Fig. 6.**
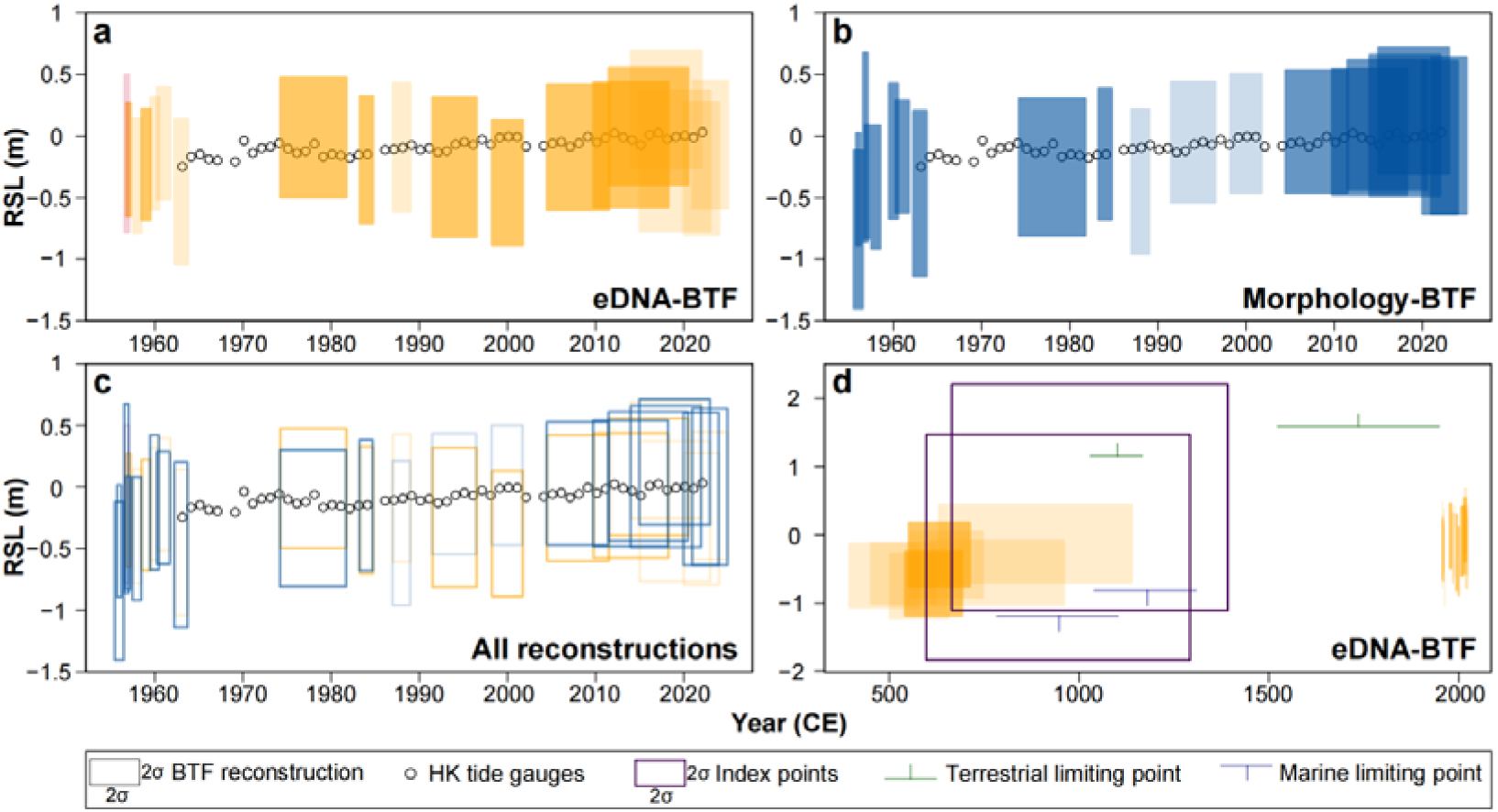
Comparison of reconstructed relative sea level (RSL) with tidal gauge records. Comparison of RSL reconstructions from foraminiferal eDNA- (a) and morphology-(b) Bayesian transfer functions (BTFs) with tide gauge data from Hong Kong. Boxes represent ±2σ vertical uncertainties and ±2σ age uncertainties for the reconstructions. RSL reconstructed from core samples with poor modern analogue were shown with transparent colors. RSL estimated by the core sample that has poor fit with tidal elevation determined by goodness-of-fit were shown by red box. Comparison of eDNA- and morphology-BTF reconstructed RSL with tidal gauge records is shown in (c). Comparison of RSL estimated by eDNA-BTF with Hong Kong tide gauge records and regional index points—delimiting the position of RSL—and limiting points— providing upper (terrestrial) or lower (marine) bounds on RSL^100^–from the Pearl River Delta is shown in (d).

## 3. Discussion

### 3.1 Influence of elevation on foraminiferal eDNA and morphological assemblages and the development of transfer functions

The reconstruction of RSL using transfer functions assumes that tidal elevation, a surrogate variable for the frequency and duration of tidal inundation, is an important ecological determinant of taxa distribution, or linearly related to another ecologically significant factor^44,45^. Indeed, elevation consistently influenced both eDNA and morphological assemblages, with samples from all three transects clustering by elevation in NMDS space (Fig. 3c, 3d). However, additional environmental factors, particularly salinity, also shaped eDNA assemblages. Salinity, which decreases with increasing elevation, was identified as a primary control on foraminiferal eDNA composition, in agreement with previous studies of estuarine foraminiferal eDNA^25^. Certain undetermined OTUs and monothalamids, for example OTU29, were strongly associated with high-salinity environments (Fig. 3b), which helps explain why elevation accounted for less variation in eDNA assemblages compared to morphological data. In contrast, tidal elevation remained the dominant variable explaining morphological assemblages (Supplementary Table 6), consistent with previous research^46–48^.

The divergence in dominant environmental controls likely originates from differences in the populations represented by the eDNA and morphological assemblages. Morphological assemblages used to develop transfer functions are typically derived from tests of dead benthic adult foraminifera to minimize temporal bias^49^. In contrast, eDNA inevitably integrates DNA from multiple sources, including living adults, juveniles, propagules, allochthonous DNA, and DNA preserved in dead foraminiferal tests^25,50,51^. For example, family Nummulitidae, despite being mineral-shelled, was identified only in the eDNA assemblage, suggesting the presence of propagules rather than adult individuals. The mobility of eDNA, similar to diatom assemblages in intertidal zones^52–54^, allows it to capture signals from both autochthonous and allochthonous taxa (e.g., planktonic taxa Globigerinitidae; Supplementary Table 4). In Mai Po and Sha Kiu mudflat samples, the low proportion of stained (live) foraminiferal tests (Supplementary Table 11) suggests limited *in situ* living populations, implying a greater contribution of propagules and allochthonous DNA to the eDNA assemblage. The strong influence of δ^13^C–indicative of a marine or terrestrial source of sediment–on the eDNA assemblage further underscores its multi-source nature. High δ^13^C values suggest greater contributions from marine organic matter sources^42^ and hence stronger contributions of allochthonous eDNA in these lower-elevation environments more open to external influences^55^.

Another factor contributing to the observed discrepancies is the inclusion of monothalamous taxa, often overlooked in morphological analyses due to their fragile tests. Monothalamids, which are abundant in eDNA assemblages from coastal and deep-sea settings^25,26,56^, may be better represented in eDNA datasets because DNA from their soft tests is more easily extracted^57^ and the short DNA barcodes of some taxa (e.g., saccamminids) improve PCR amplification and sequencing efficiency^58^. As shown in a previous eDNA-based study^25^, monothalamous taxa are sensitive to short-term physiochemical environmental changes. Their inclusion in foraminiferal eDNA assemblages likely amplifies the influence of environmental variables like salinity. Nevertheless, tidal elevation is a key ecological driver of foraminiferal zonation and exhibits strong correlations with other significant environmental variables that influence eDNA assemblage composition. Meeting this key assumption permitted the development of BTFs.

The lower uncertainty and bias of the eDNA-BTF compared to the morphology-BTF (Table 1) underscore its potential for precise and accurate elevation predictions in intertidal environments. While the morphology-BTF exhibited systematic overprediction in mangrove environments (170-200 SWLI) (Fig. 5a), likely due to the linear increase in *Miliammina* spp. abundance in higher elevations (Fig. 5c), the eDNA-BTF showed no discernible pattern in the residuals. However, relatively greater uncertainties in lower mudflat environments (<MTL) (Fig. 5a) suggests that highly-mobile propagules and allochthonous eDNA^25^ may weaken the mudflat elevation signal. This pattern aligns with findings from diatom-based studies, where low-elevation assemblages are often influenced by planktonic taxa and allochthonous sources^53,59–61^. Consistent with morphological studies^43,62,63^, the clearest elevation signals—and thus the most reliable predictions—are observed in higher-elevation mangrove assemblages, where tidal elevation exerts a more dominant ecological control.

### 3.2 Taphonomy, preservation, and application of foraminiferal eDNA for RSL reconstruction

The sedaDNA assemblage showed generally good agreement with depositional environments inferred from LDA predictions across lithostratigraphic unit (Fig. 4). Basal Unit III contained mudflat assemblages with low Miliamminidae abundance and mudflat-specific taxa (e.g., undetermined OTU29), while Units I and II showed mangrove assemblages indicated by high abundance of Miliamminidae (Fig. 4). However, the core sedaDNA assemblage had more samples (16/28) falling outside the dissimilarity threshold than the morphological assemblage (3/20; Fig. 4). For the morphological assemblage, poor analogues mainly resulted from high abundances of *Ammoastuta inepta* and *Ammonia convexa*—rare in modern environments, likely due to anthropogenic impacts^43^. Mangrove management policies at Mai Po (e.g., trimming of seedlings)^43^ may have altered modern mudflat assemblages, creating poor analogues for both morphological and sedaDNA assemblages, especially pre-1965 CE core samples (Fig. 4). In addition to human influences, poor sedaDNA analogues may reflect: (1) high taxonomic diversity, (2) eDNA’s sensitivity to subtle assemblage changes, (3) potential influence from infaunal populations, or (4) taphonomy and molecular preservation of sedaDNA.

First, the high taxonomic diversity of eDNA/sedaDNA assemblages increases heterogeneity and the proportion of rare taxa compared to morphological assemblages^64,65^, raising the likelihood of encountering taxa underrepresented in modern training sets^43^, similar to other sea-level proxies like diatoms^64,66^. For example, OTU87, primarily found in mudflats, reached 14% abundance in a core sample (1.15–1.16 m PD, Unit III), but never exceeded 5% in modern samples (Supplementary Table 4).

Second, an abrupt increase of Saccamminidae at 1.85–1.86 m PD, coinciding with the lithostratigraphic boundary between Unit I and II (Fig. 4), also resulted in a poor modern analogue. This shift may reflect changes in hydrodynamics, productivity, or other environmental factors during facies transitions^67,68^, similar to the pattern observed at the mangrove-terrestrial transition in transect MP_B (Fig. 2). Monothalamous foraminifera like Saccamminidae are highly adaptive to fluctuating conditions and can respond rapidly to environmental changes, including seasonal cycles^69^. Temporal variation in living populations may further contribute to discrepancies with modern analogues, highlighting the need for research on the seasonal dynamics of foraminiferal eDNA^70–72^.

Third, eDNA assemblages in the bioturbated mixed zone (>2.15 m PD) may have been influenced by living infaunal populations such as OTU11 and Elphidiidae, which lack preserved morphological counterparts either because they do not possess hard shells or are represented by propagule DNA^40^. Similar patterns have been reported in deep-sea sediments^36,73^, where the more abundant and pristine DNA from living foraminifera dominated the bioturbated zone^39^. Because DNA from living cells decays rapidly after cell death—primarily through endogenous nuclease activity (i.e., breakdown of nucleic acids originating from within the cell) rather than microbial activities^74–76^—eDNA below the bioturbated zone primarily represents extracellular sources and the total population flux^36^.

Lastly, both sedaDNA and morphological assemblages were affected by post-depositional processes. Below 1.31 m PD, foraminiferal tests were absent—likely due to dissolution under low pH conditions^19,77^—while sedaDNA remained preserved (Fig. 4), suggesting greater long-term stability for DNA than foraminiferal tests in such environments. The sedaDNA adsorption onto clay minerals shields genetic material from chemical and biological degradation^31,32,78^, whereas foraminiferal tests are more susceptible to dissolution and disintegration. However, a statistically significant decrease in α-diversity was observed in core samples with increasing depth (Supplementary Table 9), indicating that the sedaDNA assemblage was also affected by taphonomic processes. Low TOC (<1.3%) and lack of tests suggest reduced sedimentary shelter and protection for sedaDNA^31,36,74^. Monothalamous taxa with short barcodes (e.g., Saccamminidae) are more resilient to DNA fragmentation^58^, which may increase their apparent abundance in degraded samples and potentially lead to overestimation of PME, resulting in offsets between BTF and LDA estimates in core samples below 1.21 m PD (Fig. 4) and underprediction of RSL (Fig. 6d). While post-mortem damage and test loss preclude morphological analysis, eDNA metabarcoding can recover information from fragmented sequences as long as amplifiable DNA persists^34,36,79^, making the eDNA method more applicable under extreme conditions (e.g. low pH) where morphological approaches are limited^19,77,80–82^.

Foraminiferal sedaDNA and tests were absent, however, from a short interval between 1.25–1.31 m PD, likely resulting from a hiatus in sedimentation. This interpretation is supported by shifts in radiocarbon-determined ages and sedimentation rates at 1.31 m PD (Fig. 4). This hiatus, occurring around 800 years ago, is attributed to a lack of accommodation space^83^ caused by late Holocene RSL fall^42,84^, during which sediment accumulation ceased and sedimentary eDNA was exposed to bioturbation and weathering (prolonged warm, oxidative, and moist conditions in these subtropical wetlands) for at least 250 years, leading to the complete degradation or severe fragmentation of DNA^39,74,85–87^. The subsequent post-industrial RSL rise created accommodation for sediment deposition^88,89^, resulting in the rapid sedimentation rate (average >1.5 cm/yr) above 1.3 m PD (Supplementary Fig. 4).

Importantly, the presence of core samples with poor modern analogues does not necessarily compromise reconstruction reliability, particularly for heterogeneous datasets with high diversity^44^. Applying the eDNA-BTF to reconstruct RSL from the core showed strong agreement between the tide gauge and other geologic RSL records from the PRD (Fig. 6), even for samples lacking close modern analogs (Fig. 4), supporting the approach’s validity^90,91^. Our eDNA-based approach generated high-resolution RSL reconstructions with decadal temporal and decimeter vertical precision for two distinct chronological intervals (2σ range: 290–1703 CE and 1956 CE–present) and improved resolution for the period 290-1703 CE compared to existing geologic RSL records (Fig. 6d). Superior eDNA preservation enabled RSL reconstruction as far back as 290 CE, whereas morphological analyses could not be applied to sediments older than 1956 CE. The eDNA-BTF reconstruction demonstrated slightly lower accuracy in RSL estimation (MSE of 0.027 m² excluding samples with poor analogues) compared to the morphology-BTF (0.016 m²) and a morphology-BTF established from a salt-marsh core in New Jersey (0.014 m²)^92^ but higher precision (1σ uncertainty: 12 vs. 18 SWLI). These results demonstrate that foraminiferal eDNA is a robust proxy for reconstructing late Holocene RSL, especially where morphological methods are not feasible.

Some limitations observed, such as the impact of sedimentary hiatuses, reflect broader challenges in developing high-resolution far-field RSL records^43,48,81^ rather than inherent shortcomings of the eDNA proxy itself. The eDNA approach may have greater potential at sites characterized by rising late Holocene RSL, including mangrove archives from the Caribbean^93–95^ and mid-Pacific^96^, or salt-marsh archives from across the north Atlantic^64,90,97–99^, where more continuous sedimentary records are preserved.

### 3.3 Comparison of eDNA and morphological approaches and recommendations for future applications

The comparison between eDNA and morphological assemblages reveals critical differences influencing their utility as proxies for reconstructing past RSL. The eDNA assemblages incorporate DNA from propagules and juvenile stages, providing a broader genetic signal than morphological methods and offering valuable ecological insights. However, this broader signal introduces complexity when interpreting correlations with tidal elevation, as the detected DNA may not originate from in situ communities. To disentangle the influence of propagules, future studies could employ methods such as 10-g extraction kits^24^ or focus on the >63 µm sediment fraction by sieving sediment prior to extraction to isolate adult populations^101^.

A notable property of eDNA methods is the inclusion of undetermined OTUs within assemblages. In this study, 53% of OTUs and 37% of sequences remained undetermined, a proportion consistent with previous foraminiferal eDNA research^23,26,102^. These undetermined OTUs likely represent species whose genetic information is not yet documented in reference databases, constraining the ability to interpret their ecological preferences and hindering cross-study comparisons. Without comprehensive taxonomic identification, it is challenging to relate OTUs across molecular and morphological studies, limiting the broader application and interpretability of eDNA-based reconstructions and their integration with morphological approaches. Although these undetermined OTUs have demonstrated the ability to indicate tidal elevation, expanding sequencing efforts for foraminifera— particularly from low-energy intertidal environments—remains essential for enabling the universal application of eDNA as a robust paleoenvironmental tool.

Finally, eDNA assemblages exhibit significantly higher diversity than morphological assemblages, with 824 OTUs identified, representing both species- and subspecies-level diversity, compared to 24 morphospecies (Supplementary Table 4). An additional key advantage of eDNA is its ability to detect taxa that are often underrepresented in traditional studies^38,39^, such as monothalamids—the inclusion of which has been shown to enhance BTF performance (Supplementary Table 12). This ability to incorporate and detect genetic diversity at such high resolution could enhance the predictive power of eDNA in paleoenvironmental applications, particularly if this diversity reflects ecological sensitivity to environmental variables.

## 4. Methodology

### 4.1 Site selection

The study area is situated in southeast Deep Bay on the eastern Pearl River Estuary (Fig. 1b). This subtropical region experiences an average maximum temperature of ∼30 °C in July and minimum of 17 °C in January. Average annual precipitation totals 2,396 mm (2019-2023), and most rainfall is recorded from June to September^103^. Brackish salinities (9.7-29.7) are observed in Deep Bay^104^ due to the discharge of the Pearl River and nearby Shenzhen River, and its average water depth is ∼2.9 m^105^. Tides display a semidiurnal pattern in this area, with an amplitude of 1.96 m (MHHW-MLLW) recorded by the nearest tide gauge station at Tsim Bei Tsui (Fig. 1c).

Two study sites, Mai Po and Sha Kiu, were chosen to conduct sampling of modern sediment samples. The Mai Po Nature Reserve, which encompasses a large area of intertidal mudflat and mangrove, has been under management of the World Wide Fund for Nature (WWF) since 1975. Mangrove flora in Mai Po are dominated by *Kandelia candel*, *Aegiceras corniculatum,* and *Avicennia marina*^106^. We chose Mai Po to conduct modern sample collection because the long-term management of this site ensures minimal direct human influence. The sediment core (MPSC01) was collected from the upper-mangrove of Mai Po (Fig. 1c) due to the long-standing presence of mangrove forest since at least the mid-20th century^42^. Sha Kiu, an area encompassing mudflat and mangrove environments, with flora dominated by *Aegicera corniculatum* and *Sonneratia apetala*, was also selected for modern sample collection due to its distinctive transition from mudflat to mangrove habitats, which we considered experienced relatively less human activity compared to other areas along the Deep Bay coast.

### 4.2 Sample collection

To investigate the vertical zonation of foraminiferal environmental DNA (eDNA) assemblages and to construct the modern training set for sea-level reconstruction, a total of 60 surface samples were collected along three transects: two in Mai Po (MP_A and MP_B) and one in Sha Kiu (SK), covering the subtidal, mudflat, mangrove and terrestrial forest areas (Fig. 1c). To evenly sample the elevation and environmental gradient, sampling stations were spaced at 5-cm elevation intervals, and in the flat mangrove forest of MP_A, every 30 m in distance. At each sampling station, surface (1-cm deep) sediment samples were collected individually for analysis of eDNA, morphology, and other environmental variables from a 10-cm^2^ area. Individual disposable spatulas were used to place samples for eDNA analysis in 50 mL Falcon centrifuge tubes. Sediment samples from the subtidal zone were collected using a Van Veen grab sampler^107^, from which ∼2 cm^3^ of the top 1 cm of surface sediments were sampled using spatulas and stored in 15 mL Falcon tubes. To minimize cross-contamination, sample collection was conducted with disposable sterile gloves, and all non-disposable equipment used in sample collection (e.g., spatulas, Van Veen grab sampler) were sterilized with 10% bleach prior to use and between each sampling station. All eDNA samples were kept on ice in the field prior to storage in a −20 °C freezer back in the laboratory on the sampling day^108^. Samples for traditional morphological analysis were preserved in an ethanol-calcite buffer solution and stained using rose Bengal in the field to differentiate between living and dead foraminifera^109^. Both samples for morphology and analysis of environmental variables were also kept in a cool box on ice in the field and stored in a 4 °C refrigerator in the laboratory.

Two 290-cm long replicate sediment cores (MPSC01) were collected from the Mai Po mangrove using an Eijkelkamp peat sampler in overlapping 50-cm intervals to minimize influence of compaction and contamination (Fig. 1c). Cores were sealed with plastic wrap in PVC pipes and kept on ice before returning to the laboratory. For the collection of the first replicate core used for sedimentary ancient DNA (sedaDNA) analysis, all materials and equipment (e.g., Eijkelkamp peat sampler, PVC pipes) were sterilized with 10% bleach before each 50-cm interval was sampled to prevent cross-contamination. Subsampling–conducted in a sterilized laboratory–was performed every 5 cm downcore using a disposable spatula. ∼1 cm^3^ of sediment was sampled from the inner part of the core to further minimize contamination^110^ (because the sample’s exterior surfaces came into contact with the peat sampler and PVC pipe during coring and storage) and stored in 15 ml Falcon tubes in a −20 [freezer. The second core replicate (collected adjacent to the first) was subsampled for morphological, geochemical and chronological analyses. For morphological analysis, samples from a 1-cm thick interval from the second replicate core were placed into 50 ml centrifuge tubes at the same sampling interval as sedaDNA, and preserved in an ethanol-calcite buffer solution and stored at 4 [on the sampling day.

To construct an in-house database of foraminiferal DNA, surface (top 1 cm) sediment samples were collected from the upper-mangrove, lower-mangrove and upper-mudflat environments along MP_A transect and from the mangrove fringe environment along transect SK (Fig. 1c; Supplementary Fig. 1). Samples were collected following the same sampling protocol as surface samples for eDNA metabarcoding. Collected samples were temporarily stored in a cool box with ice and stored in a 4 °C refrigerator in the laboratory on the sampling day.

The elevation of surface samples and the core top were measured and related to a temporary benchmark using an auto level. The benchmark elevation was measured relative to the Hong Kong Principal Datum (PD) using a Leica GS18 GNSS system. Given the reduced tidal amplitude recorded by the Solinst Levelogger® 5 at the MP_B transect^43^, elevation of each surface sampling station and core top was converted to standardized water level index units (SWLI)^100^, where a value of 100 represents MTL and a value of 200 represents MHHW, based on observations from the Tsim Bei Tsui tide gauge (Fig. 1c). The location of each sampling station was recorded using a handheld GPS.

### 4.3 Single-cell foraminifera DNA isolation and Sanger sequencing

To construct an in-house database of foraminiferal DNA to complement the current public database, we conducted DNA barcoding with single individuals of foraminifera. For isolating single individuals, sediment samples were wet-sieved (63-500 μm) using artificial sea water (ASW, Red Sea Salt, salinity = 34) within a week of collection^24^. The 63-500 μm size fraction, the same size fraction that we used for morphological analysis of foraminifera, was examined in ASW under a stereomicroscope for signs of vitality. Foraminifera exhibiting pigmented cytoplasm within the test and an empty last chamber were considered to be viable and placed with a bleach-sterilized brush into a picking dish with ASW for vitality assessment for maximum of 8 hours^108,111^. Active foraminifera (exhibiting reticulopodial activity and movement) were cleaned and photographed using a Leica stereomicroscope for morphological identification. After imaging, samples were placed individually into 50 μl DOC (Deoxycholate) buffer, smashed using a sterile disposable crusher and then incubated at 60 °C for an hour^112^. PCR amplification was conducted using foraminifera-specific primers s14F1-sB and s14F1-N6^112,113^ following protocol described in Darling et al. (2016)^113^. For individuals that did not show any visible bands in the agarose gel electrophoresis of the nested PCR, primers s14F1-s17^112^ were used in a 2^nd^ nested PCR amplification instead of s14F1-N6. Successfully amplified samples were sequenced with the Sanger method (BGI, Guangzhou). Sequences from 13 foraminiferal individuals were successfully obtained by single-individual DNA barcoding and used to construct the in-house database. Four new *Ammonia* phylotypes and one new *Cribroelphidium* phylotype were identified (Supplementary Table 3). The full sequences were uploaded to the NCBI Genbank database.

### 4.4 eDNA/sedaDNA analysis

#### 4.4.1 eDNA/sedaDNA extraction, amplification, and sequencing

All eDNA/sedaDNA laboratory procedures were performed in a molecular laboratory that had no prior history of foraminiferal DNA research. All DNA extraction, PCR, and library construction procedures were conducted within a safety cabinet, with all materials pre-exposed to UV irradiation for ≥15 minutes^114^. Except for disposable sterile or pre-autoclaved equipment, all reusable materials (e.g., pipettes) were sterilized with 70% ethanol and/or 10% bleach before use and between sample processing steps to minimize cross-contamination. DNA was extracted from surface sediment and core samples using the 0.5-g PowerSoil® DNA Isolation Kit (QIAGEN) in a sterile laboratory environment following the manufacturer protocol, and stored in a −80 [freezer. DNA extraction was conducted in duplicate (2 technical replicates) for all surface and core samples to maximize genetic diversity^25^. Specifically, core sample extractions were performed from the deepest to shallowest layers to minimize potential cross-contamination from upper, typically higher-concentration sediment layers. We performed the polymerase chain reaction (PCR) by amplifying the specific hypervariable region 37f of foraminiferal 18s SSU rDNA using primers s14F1 and s15^102^. To multiplex PCR products for sequencing, both primers were synthesized with the overhang adapter sequences for compatibility with Illumina index and sequencing adapters following the Illumina MiSeq System sequencing protocol: 5’ Adaptor - s14F1 3’ and 5’ Adaptor – s15 3’. For each extracted sample, PCR was conducted in duplicate (2 technical replicates) to minimize potential PCR biases^115^. Positive (DNA using synthetic *Miliammina fusca* clone 4 small subunit ribosomal RNA gene as template, gBlocks®, 787 bp) and negative (blank) controls were applied during PCR amplification and library construction to indicate positive amplification and to detect possible cross-contamination. The synthetic sequence used as a positive control was artificially modified during synthesis through nucleotide substitutions in our target barcode regions to enable contamination detection. PCR mix was prepared in 25 µl reaction, containing 2.5 μl of samples, 12.5 μl of Taq PCR Master Mix (QIAGEN), 4 μl of each primer (1 µM), 1.25 μl of Mg^2+^ (1 µM) and 0.75 μl of Millq water. PCR was performed as follows: an initial denaturation at 94°C for 3 minutes, followed by 45 cycles of 94 °C for 30 s, 55 °C for 30s, and 72 °C for 45s, with a final extension at 72 °C for 5 minutes. The amplified products underwent a first clean-up using the AMPpure XP beads (Beckman Coulter, Singapore). A 20 μl aliquot of technical PCR replicates were pooled before first clean-up. Index PCR was then performed on the cleaned products to attach dual indices on Illumina sequencing adapters using the Nextera XT Index. The reaction mix comprises 25 µl 2× KAPA HiFi HotStart ReadyMix, 5 µl of each index primer (10 µM), 10 µl of PCR grade water, and 5 µL cleaned PCR product. Index PCR amplification condition was set up as follows: initial denaturation at 95 °C for 3 minutes, 8 cycles of denaturation for 30 s at 95 °C, annealing for 30 s at 55 °C and extension for 30 s at 72 °C and a final extension for 5 min at 72 °C. The 2^nd^ PCR clean-up was conducted consecutively, and in total 177 cleaned foraminiferal eDNA/sedaDNA samples (including negative control, even when no visible bands were observed during agarose gel electrophoresis of the PCR products) were then pooled to 20 μM using 10 mM Tris (pH 8.5) and sent to Novogene Co, Ltd for sequencing on the NovaSeq System PE 150 platform. To minimize potential cross-contamination of the sedaDNA from modern eDNA assemblage, all laboratory procedures before the second PCR clean-up were performed separately for modern and core samples.

#### 4.4.2 Bioinformatics

Raw sequence data were demultiplexed, and primer sequences were removed by Cutadapt v2.8^116^. The resulting pair-end reads were then merged by PEAR v0.9.11 with a quality threshold of 26^117^. The concatenated sequence file underwent subsequent denoising using VSEARCH-UNOISE3 with default alpha diversity parameters and a minimum size threshold of 4, followed by chimera removal with VSEARCH-UCHIME^118^. Sequences with length shorter than 90 bp and longer than 230 bp were discarded using VSEARCH. Sequences exhibiting >97% similarity were clustered into operational taxonomic units (OTUs)^119^ by VSEARCH^118^, and a resulting OTU table (numbers of OTUs per sample) containing 36,144,049 sequence number was constructed using VSEARCH-USEARCH. Quality control was applied to discard OTUs that: (1) had fewer than 10 reads^102^, or (2) appeared in negative controls^120^ with read counts equal to or exceeding those in any sample^114^, to minimize index hopping artifacts^121^. Technical extraction replicates from the same surface or core sample were pooled prior to subsequent processing. Taxonomic assignments were made using BLASTn v2.12.0^122^ with e-value of 0.001 referencing the Genbank nucleotide (nt) database (downloaded on 13 September 2024) from the National Center for Biotechnology Information (NCBI)^123^, following previous foraminiferal eDNA studies^25,36^. The in-house database was also used as a complementary reference^25^. The python script taxonomy_assignment_BLAST_V1.py^124^ was used for processing the BLAST results to assign final taxonomic classifications to the species, family, and phylum level based on identity thresholds of 98%, 90%^125^, and 80%^126^ respectively, utilizing the top BLAST hits and best consensus taxonomy that met the predefined identity percentage thresholds. Compared to assigning OTUs to species, genus, family, and phylum levels at 99%, 95%, 90%, and 80% identity thresholds, respectively, this approach demonstrated superior performance in relative sea-level (RSL) reconstruction (Supplementary Table 12). Where BLAST results showed conflicting taxonomic assignments, OTUs were assigned at higher taxonomic ranks (genus or order level) to establish consensus identification^114^. Where taxonomic assignments differed between GenBank and in-house databases, we adopted the in-house database classification^114^. OTUs taxonomic assignments were conducted referring to the NCBI Taxonomy database, followed by manual verification and correction against taxonomic records in the World Register of Marine Species (WoRMS)^127^. OTUs with identities between 80 to 90% thresholds (assigned to Foraminifera phylum) and OTUs that were assigned to an unknown taxonomy level of a foraminifera sequence were classified as “undetermined OTUs” of foraminifera^126^ and were treated as individual taxa in the subsequent analysis. OTUs with identity thresholds <80% were not considered to be foraminifera and were removed^24^.

Although clustering sequences into amplicon sequence variants (ASVs) has become increasingly popular due to its ability to retain more genetic diversity^128^, we did not use this approach. This decision was based on the large proportion of undetermined sequences in our dataset and our use of individual undetermined sequences for the analysis. Introducing higher taxonomic diversity would involve taxa with low fractional abundance, which could reduce statistical reliability in RSL reconstruction given the fixed total count^129–131^.

### 4.5 Environmental variables and ***δ***^13^C and C/N geochemistry

To assess influence of environmental parameters on foraminiferal eDNA, a suite of environmental variables was measured from modern samples: salinity, pH, and grain size. Sediment samples were centrifuged at 3000 rpm for 30 minutes to measure porewater salinity and pH using a Thermo Scientific™ A3255 pH/conductivity multimeter^132^. For samples that yielded insufficient porewater for measurement, 5 mL of Milli-Q water was introduced to the samples before centrifugation and measurement^59^. Sediment samples for grain size measurement were treated with 30% hydrogen peroxide to remove organic carbon and then dispersed using Calgon solution. Grain size analysis was performed using a Beckman-Coulter LS 13 320 laser particle size analyzer. The software program GRADISTAT 9.1 was used to calculate grain size statistics^133^.

To evaluate factors influencing the composition of modern foraminiferal eDNA assemblages and to provide constraints for RSL reconstructions, stable carbon isotope geochemistry parameters—including total organic carbon (TOC), total nitrogen (TN), and δ^13^C—were measured in both modern and core samples. Prior to δ^13^C, TOC, and TN analysis, sediment samples were pretreated with 5% hydrochloric acid (HCl) for 24 hours to remove carbonate material, followed by rinsing with ∼1500 ml Milli-Q water^43^. δ^13^C, TOC, and TN were measured at the Stable Isotope Laboratory at the University of Hong Kong using a EuroVector EA3028 Nu Horizon isotope-ratio mass spectrometer (IRMS) and a EA Isolink Elemental Analyser. δ^13^C values were normalized to the Vienna Pee Dee Belemnite (VPDB) scale with reference to triplicate analysis of certified reference materials supplied by the United States Geological Survey, USGS40 (average ^15^N = −4.52 and ^13^C = −26.39) and USGS41a (average ^15^N = 47.55 and ^13^C = 36.55).

### 4.6 Morphological foraminiferal analysis

To provide context for foraminiferal assemblages inferred through eDNA and sedaDNA analyses, traditional morphological analysis was performed on surface and core sediment samples. The 63-500 µm sediment fraction was isolated through wet-sieving prior to wet-picking of foraminiferal tests under a stereomicroscope^134,135^. Where possible, a minimum of 100 individuals were enumerated from each sample to ensure robust transfer function development^131,136^. Taxonomy followed earlier work from the study site^43^ (Supplementary Table 13). Poor specimen preservation prevented species-level identification in several genera. Therefore, species of *Ammobaculites* (*A. exiguous, A. agglutinans*), *Miliammina* (*M. fusca, M. obliqua, M. petila*), and two *Ammonia* species (*A. beccarii, A. confertitesta*) were combined under *Ammobaculites* spp., *Miliammina* spp., and *Ammonia* spp., respectively^47,80,93,137^. Further attempts to combine *A. tepida* within *Ammonia* spp. to avoid misidentification resulted in anomalies in PME prediction (Supplementary Fig. 5), and were therefore abandoned. The morphological dataset comprises only the dead assemblage (unstained by rose Bengal) to avoid the influence of temporally variable living foraminiferal populations^49^. To minimize bias introduced by rare taxa, only the taxa with relative abundance >5% in at least one surface or core subsample were included in the analysis^43^.

### 4.7 ^14^C dating

The chronology of core MPSC01 was constructed from radiocarbon-dated plant fragments and fine-fraction organic mud from 15 core samples (Supplementary Table 14). Plant fragments, including mangrove bark, branches and twigs deemed to be deposited horizontally on the former sediment surface with no clear evidence of reworking, were chosen as dating materials^21,97^. Plant fragments were examined and cleaned with Milli-Q water under a stereomicroscope to remove all visible rootlets and adhering material. Fine-fraction organic mud samples for radiocarbon dating were prepared by isolating the <63-µm fraction on a 0.47 µm quartz fiber filter under filtration, followed by oven drying for 24 hours at 40 °C. Dried organic mud samples were weighed, wrapped in aluminum foil, placed in clean glass jars, and stored in a desiccator until further processing. To minimize modern carbon contamination and cross-contamination, all equipment was pre-treated before use. Quartz fiber filters and aluminum foil were baked at 550 °C for 4 hours, then cooled in a desiccator. All sieves and processing tools were cleaned between samples using a four-step protocol: (1) washing with soapy water; (2) rinsing with tap water; (3) soaking in 10% HCl for ∼30 seconds; and (4) triple rinsing with Milli-Q water. Additionally, the 63-µm sieve was sonicated for 15 minutes before processing each sample. Samples were submitted either to Beta Analytic in Miami, FL, USA, using accelerator mass spectrometry (AMS) dating, or to the Yale Analytical and Stable Isotope Center (YASIC) in New Haven, CT, USA, using the MICADAS AMS system.

An age model for core MPSC01 was constructed using the Bacon age-depth model^138^, incorporating all radiocarbon-dated samples to estimate the age with 2σ uncertainty of each eDNA/morphological core sample (Supplementary Fig. 4; Supplementary Table 14). The Intcal20 calibration curve^139^ was used for calibrating the conventional ages to calendar years, and the post-1950 CE samples were calibrated using the bomb spike dataset^140^.

### 4.8 Statistical analysis

#### 4.8.1 Alpha diversity analysis

To estimate the α-diversity of eDNA and sedaDNA assemblages, rarefaction^141^ was performed to standardize each sample to the lowest sequencing depth of the dataset (10,591 reads), a depth at which most samples reached a plateau in rarefaction curves (Supplementary Fig. 6). Two samples (one modern and one core sample) with insufficient read counts were excluded from further analysis. For the remaining samples, three α-diversity metrics were calculated: observed OTUs^128^, Shannon index^142^ and Chao1 index^143^. Differences in α-diversity were assessed using one-way ANOVA with 1,000 permutations (p-value threshold = 0.05)^114,144^, focusing on: (1) variation among modern samples across six identified environments; (2) differences between modern and core samples; and (3) differences between core samples from Units I/II and Unit III.

Similarly, α-diversity metrics—including taxa richness, Shannon index, and Chao1 index—were calculated for the morphological assemblage, with a focus on: (1) differences among samples from six modern environments, and (2) comparisons between modern and core samples.

#### 4.8.2 Multivariate Ordination analysis

Non-Metric Multidimensional Scaling (NMDS) analysis was conducted on modern foraminiferal eDNA and morphological assemblages to highlight dissimilarities in terms of community composition between sampling stations with different elevations^145^. NMDS was performed using a Bray-Curtis dissimilarity matrix based on taxa relative abundance data. The resulting ordination plot displays the first two NMDS dimensions, where the spatial arrangement of points reflects the compositional dissimilarity among stations. A Permutational Multivariate Analysis of Variance (PERMANOVA) with 999 permutations, based on Bray-Curtis distance, was performed to assess whether the observed grouping patterns in the NMDS ordination were significantly explained by elevation differences^146^. Analysis was conducted using the “vegan” package (v2.6-4) in R (v4.3.1)^147^.

Canonical correspondence analysis (CCA) was performed with relative abundance data to examine the correlation between the contemporary foraminiferal eDNA and morphological assemblages with environmental variables across all modern samples^148^, also conducted using the “vegan” package (v2.6-4) in R^147^. For both eDNA and morphological assemblages, we included only taxa that met two criteria in the CCA: (1) >5% relative abundance in at least one modern or core sample, and (2) presence in more than one sample within the modern training set^43,46^. In CCA and partial CCA (pCCA), the degree of variation explained by environmental variables are measured by canonical eigenvalues^46,60^.

#### 4.8.3 Transfer function development

##### 4.8.3.1 Transfer function construction

A Bayesian transfer function (BTF) was constructed using modern training sets of foraminiferal eDNA and morphological assemblages. Taxa with >5% abundance of the assemblage in any sample were included in the model to minimize uncertainty associated with low-abundance taxa^92,129,130^. The BTF approach uses penalized spline functions to model non-parametric species response curves, enabling the representation of complex, multimodal, and non-Gaussian responses to elevation. Informative priors derived from elevation-dependent environments based on stable carbon isotope geochemistry^92^ were incorporated to constrain predictions.

For the morphological assemblage, counts of foraminiferal taxa served as model input, excluding the core sample at 1.55-1.56 m PD due to insufficient counts. This approach reduces uncertainty associated with larger count sizes while maintaining flexibility to capture multi-modal and non-Gaussian species responses^92^. For the eDNA assemblage, raw sequence counts were not used due to the discrepancies in sequencing depth between samples^149^. Instead, rarefied sequences—subsampled to a uniform sequencing depth^141^—were used as model input, as in the α-diversity calculation. While rarefying DNA data requires caution^150^, compared to normalization methods^151^, this approach preserves the discrete nature of the dataset and avoids violating the BTF assumption of multinomial taxa distribution^92,152^.

Ten-fold cross-validation was conducted to evaluate BTF performance for eDNA and morphological assemblages by comparing the root mean squared error of prediction (RMSEP), correlation between predicted and observed values (R^2^), average 1σ uncertainty, average residual and average absolute residual between predicted and observed values of surface samples^44^. A single-pass approach was adopted to screen out samples with residuals exceeding two standard deviations from the mean during BTF construction^153^.

Given the lack of established best practices for developing BTFs from eDNA data, sensitivity tests were conducted to assess how training set composition influenced BTF performance. These tests included (1) the use of rarefied vs. non-rarefied taxa, (2) using different abundance thresholds for removing rare taxa (e.g., 1, 2, 5%), (3) using different taxonomy assignment criteria (e.g., using 99% and 95% identity for species- and genus-level assignment), (4) inclusion vs. exclusion of undetermined OTUs, (5) inclusion vs. exclusion of soft-walled monothalamids, (6) using training sets with stronger elevation signals (e.g., mangrove-only or mangrove and mudflat subsets), and (7) the use of site-specific training sets. These tests confirmed that excluding low-abundance taxa through the application of an abundance threshold^129,130^—for which 5% was determined to be optimal in our study—mitigates the potential impact of rare sequence loss during rarefaction, ensuring robust BTF performance. Using OTUs clustered at the lowest taxonomic level allowed input taxa to represent varying taxonomic levels, while treating undetermined OTUs as individual taxa and including monothalamids further improved model performance. Additionally, the training set incorporating samples across all environments yielded the most precise and accurate RSL reconstructions (Supplementary Fig. 7; Supplementary Table 12). The eDNA-BTF also demonstrated better performance in cross-validation and higher accuracy compared to the widely used weighted-averaging partial least squares regression (WA-PLS) transfer function (Supplementary Table 15), providing an additional rationale for selecting this BTF approach.

##### 4.8.3.2 Linear Discriminant Analysis (LDA) Priors

Linear discriminant analysis (LDA) was applied to core samples from MPSC01 to estimate prior elevation ranges used in the BTF^92^. Linear discriminant functions were derived from geochemical data (δ^13^C, TOC, and C/N) collected from modern surface samples across six elevation-dependent environmental zones^43^, including subtidal, mudflat, mangrove fringe, mangrove, terrestrial transition, and terrestrial environments (Supplementary Fig. 1; Supplementary Table 1). They were then applied to core samples to estimate the likelihood of each sample belonging to the predefined environmental zones. Core samples were assigned to a single environmental zone if their probability of affiliation exceeded a threshold of >0.9. Samples with probabilities <0.9 were assigned to multiple environmental zones. The observed elevation range corresponding to each environmental zone was used as prior information in the BTF (Supplementary Table 16).

##### 4.8.3.3 Application of Bayesian Transfer Function (BTF) to Core Samples

The eDNA- and morphology-BTFs were applied to core MPSC01 to reconstruct paleo-mangrove elevation (PME). The performance and reliability of the BTF models was evaluated using: (1) 1σ uncertainty derived from core prediction results, and (2) dissimilarity and goodness-of-fit thresholds between modern and fossil assemblages and their elevation^90^ (Fig. 4).

Dissimilarity between modern and core samples was assessed using Bray-Curtis metrics^90^, consistent with the approach applied in the NMDS analysis. Thresholds of 20% dissimilarity for morphological assemblages and 10% for eDNA assemblages were applied, reflecting the higher diversity and heterogeneity of eDNA datasets^65^. Core samples exceeding these thresholds were interpreted cautiously due to the lack of modern counterparts^90^. Goodness-of-fit statistics were calculated by fitting core samples into a constrained ordination (CCA) derived from the modern training set to evaluate their fit to elevation^90^. A 95% threshold of squared residual lengths was used to identify poor fits.

### 4.9 Validation of RSL reconstructions using regional tide gauge and geologic data

RSL was calculated by subtracting the PME reconstructed by BTFs from the elevation of each core sample^64^, both expressed relative to the Principal Datum of Hong Kong^154^. Vertical uncertainties were derived from sample-specific 2σ uncertainties from the BTFs. A chronology for RSL changes was obtained from the age-depth model (Section 4.7).

Reconstructed RSL was compared with tide gauge data from Hong Kong to validate accuracy. Due to the rapid sedimentation rate observed in the upper portion of MPSC01, annual average tide gauge measurements were calculated relative to a 5-year average (2017–2022 CE) from tide gauges at Tai Po Kau, Tai Miu Wan, Tsim Bei Tsui, Shek Pik, and Quarry Bay^43^ (Supplementary Fig. 8). This comparison spans reconstructed RSL and observed measurements from 1963 to 2022 (Fig. 6a, 6b). The mean squared error (MSE) between reconstructed RSL midpoints and tide gauge observations was calculated to assess reconstruction accuracy (e.g., Cahill et al., 2016)^92^. For older samples in age than the tide gauge records, the eDNA-BTF reconstruction was evaluated by comparison with regional sea-level index points (defining RSL position and age) and terrestrial/marine limiting points (indicating upper and lower RSL limits)^100^ from the PRD^42,155^, to confirm its accuracy.

## Supporting information

Supplementary Figure - Foraminiferal environmental DNA reveals late Holocene sea-level changes

Supplementary Table - Foraminiferal environmental DNA reveals late Holocene sea-level changes

## Data availability

All supplementary tables are presented in the “Supplementary Table - Foraminiferal environmental DNA reveals late Holocene sea-level changes” excel file. All supplementary figures and corresponding legends are presented in “Supplementary Figure - Foraminiferal environmental DNA reveals late Holocene sea-level changes” file. Raw sequence data of eDNA samples this study have been deposited in National Center for Biotechnology Information (NCBI) Sequence Read Archive (SRA) under the BioProject ID <u>PRJNA1273657</u>(https://dataview.ncbi.nlm.nih.gov/object/PRJNA1273657?reviewer=jdt0fihub4idpug78ea1k1dkk9). All barcoded sequences have been deposited in NCBI Genbank with accession numbers PV764368-PV764380.

## Code availability

Codes of bioinformatic pipeline and eDNA Bayesian transfer function are available from the GitHub repository: https://github.com/FrancisLauzj-hku/Forams_metabarcoding_sea_level.

## Acknowledgements

This research was supported by the Research Grants Council of Hong Kong (GRF Project no.: 27300221 and 17303925). We would like to thank the World Wide Fund for Nature for granting us permission to conduct research within the natural reserve. We appreciate WWF’s efforts in maintaining natural habitats and conserving wildlife in Hong Kong. Our sincere thanks also go to Miss Qin Yonghui, Mr. Gao Chengcheng, and Miss Eunice Leung (Names are listed in no particular order) for their tremendous assistance with fieldwork. We gratefully acknowledge Mr. Gu Yifei for his support in conducting data analysis. Gratitude expresses to Prof. Niamh Cahill for providing insights on Bayesian transfer function application.

## Author contributions

Liu Zhaojia conducted sample collection, laboratory work, data analysis and wrote the original manuscript. Nicole S. Khan conceptualized this study, provided supervision, assisted with sample collection and laboratory work, and contributed to manuscript writing, reviewing and editing. Howard K.Y. Yu conducted sample collection and laboratory work, and reviewed the manuscript. Arthur Chung assisted with laboratory work and data analysis, and reviewed the manuscript. Magali Schweizer provided support for laboratory work and data analysis, and contributed to manuscript review and editing. Celia Schunter conceptualized the study, provided supervision and laboratory support, and contributed to manuscript reviewing and editing.

