## Supplementary Figure - Foraminiferal environmental DNA reveals late Holocene sea-level changes for "Foraminiferal environmental DNA reveals late Holocene sea-level changes"

**Other supplementary material includes:** Supplementary Tables 1 to 16 (provided in the excel file "*Supplementary Table - Foraminiferal environmental DNA reveals late Holocene sea-level changes*")

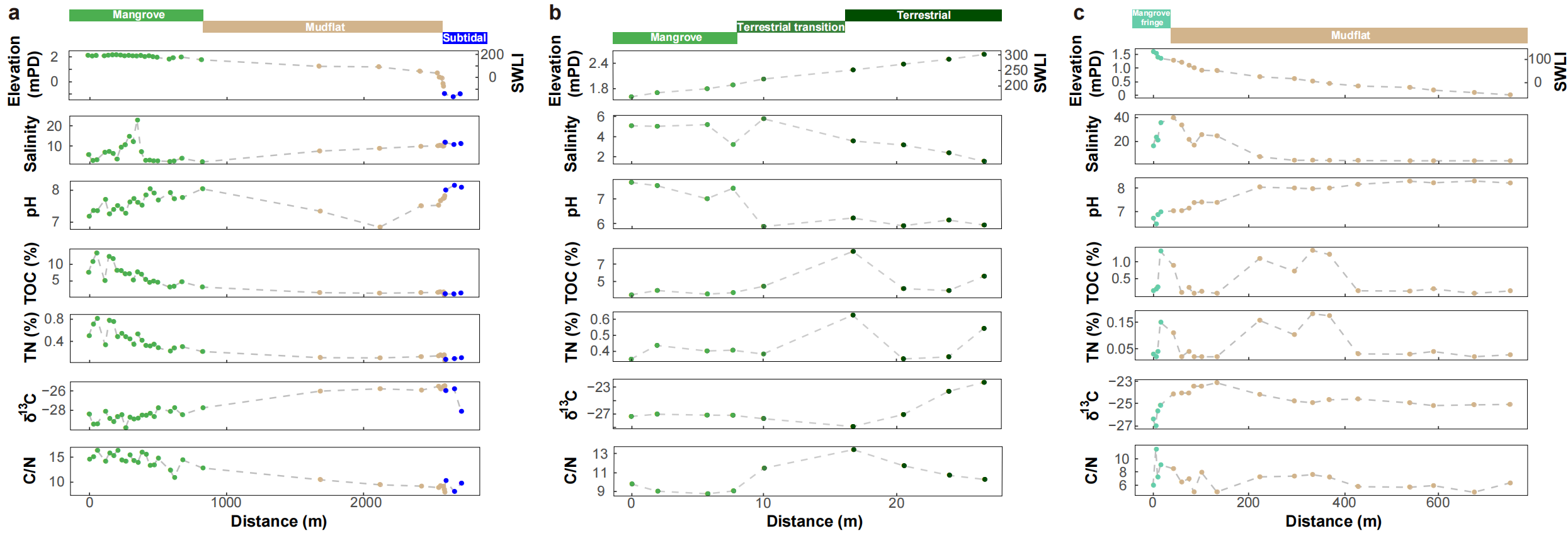
**Supplementary Fig. 1** Environmental variables measured at each sampling stations along three surface sample transects and observed environmental zone relevant to distance and elevation. (a) Mai Po A transect (MP_A). (b) Mai Po B transect (MP_B). (c) Sha Kiu transect (SK). Salinity, pH, total organic carbon content (TOC), total nitrogen (TN), δ^13^C and C/N ratio) at each sampling stations relative to distance are plotted.

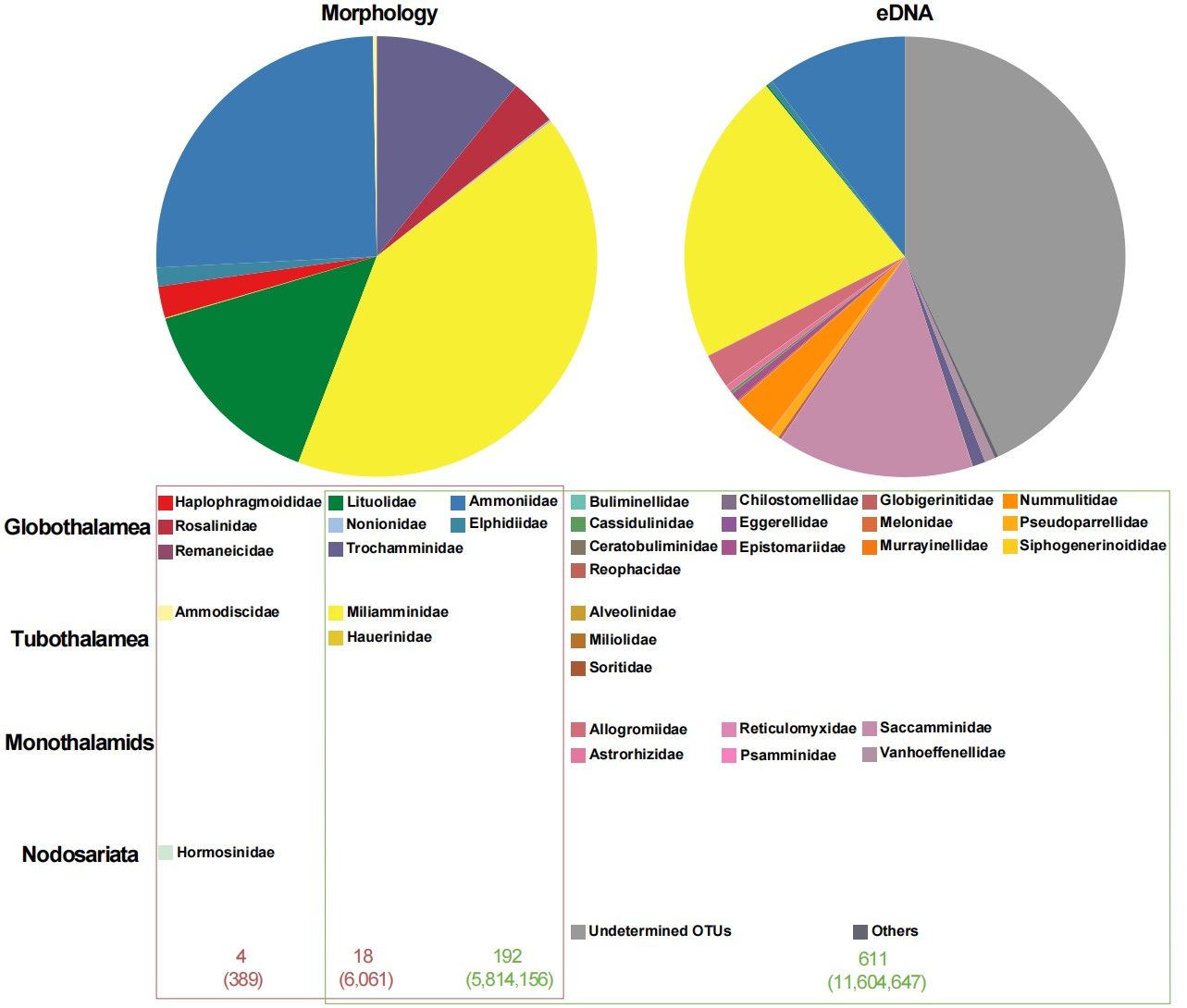

**Supplementary Fig. 2** Abundance of taxa at Family level of morphological and eDNA assemblages in modern samples. The proportion of each taxon presented in the pie chart is related to its counts/reads number in the dataset. Red numbers correspond to morphological data, indicating the number of morphospecies (with the number of specimens in brackets). Green numbers correspond to molecular data, indicating the number of OTUs (with the number of sequences in brackets). “Undetermined OTUs” with identities between 80 to 90% thresholds (assigned to Foraminifera phylum) and OTUs that were assigned to an unknown taxonomy level of a foraminifera sequence. “Others” includes OTUs assigned to the order level.

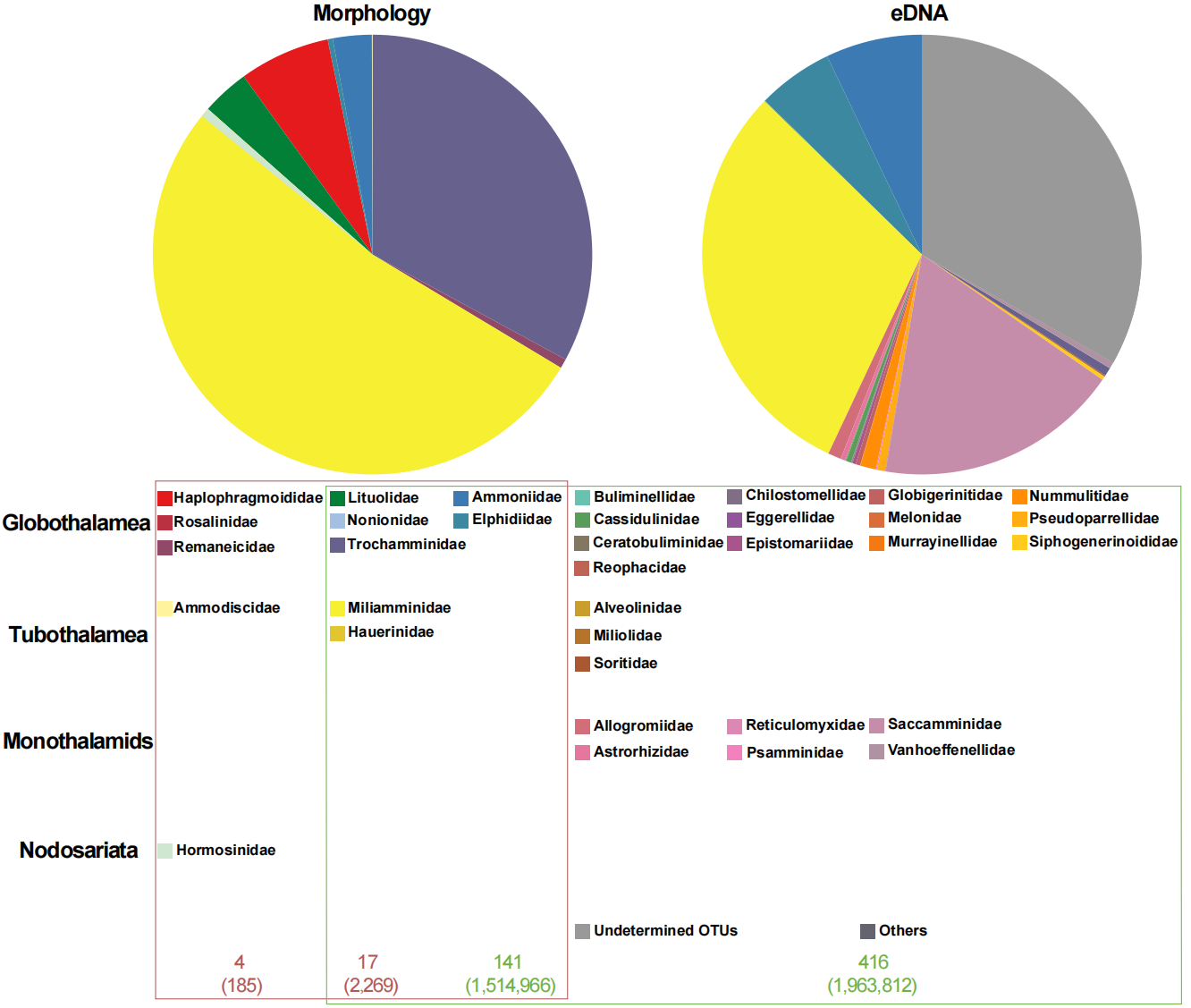

**Supplementary Fig. 3** Abundance of taxa at Family level of morphological and eDNA assemblages in core MPSC01. Counts/reads number of morphological and eDNA assemblage are shown in red/green numbers. “Others” includes OTUs assigned to the order level.

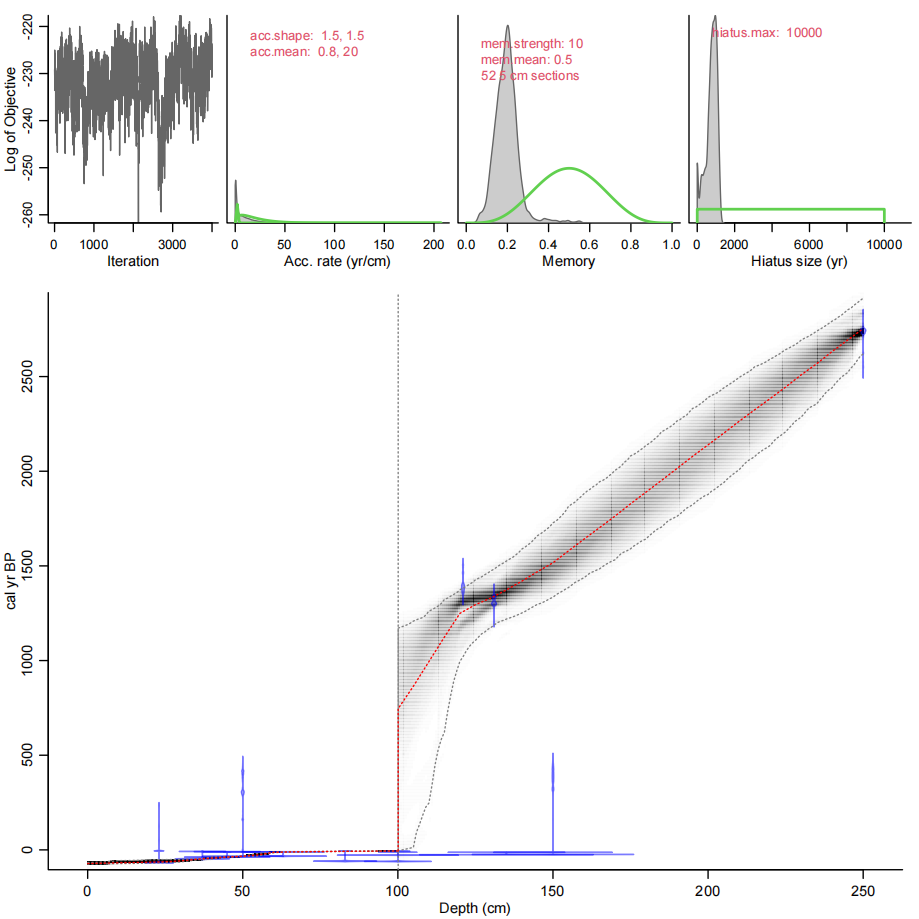

**Supplementary Fig. 4** Bacon age-depth model of core MPSC01. The upper panels show the Markov Chain Monte Carlo (MCMC) iterations on the left, where good runs display a stationary distribution with minimal variation between neighboring iterations. The middle panel presents the prior (green) and posterior (grey) distributions for the accumulation rate and its variability (memory). The bottom panel depicts the calibrated ^14^C dates (transparent blue) alongside the age-depth model, with darker grey indicating more probable calendar ages, grey stippled lines representing 95% confidence intervals, and the red curve showing the single 'best' model based on the mean age at each depth. The vertical dashed line marks the depth of the hiatus. All radiocarbon-dated samples were used to construct the age-depth model. The model captured 67% of the samples within the 2-sigma uncertainty range. For core MPSC01, the age-depth model above the 100 cm hiatus aligns the chronology reported in a previous study using the same core^43^. Below the 100 cm hiatus, the model shows a similar chronological trend to previous studies from the area^42^. Radiocarbon dates from plant material at depths of 135 cm and 150 cm, and from organic mud at 150 cm, were not captured by the model. These samples yielded anomalously young dates, likely due to disturbance.

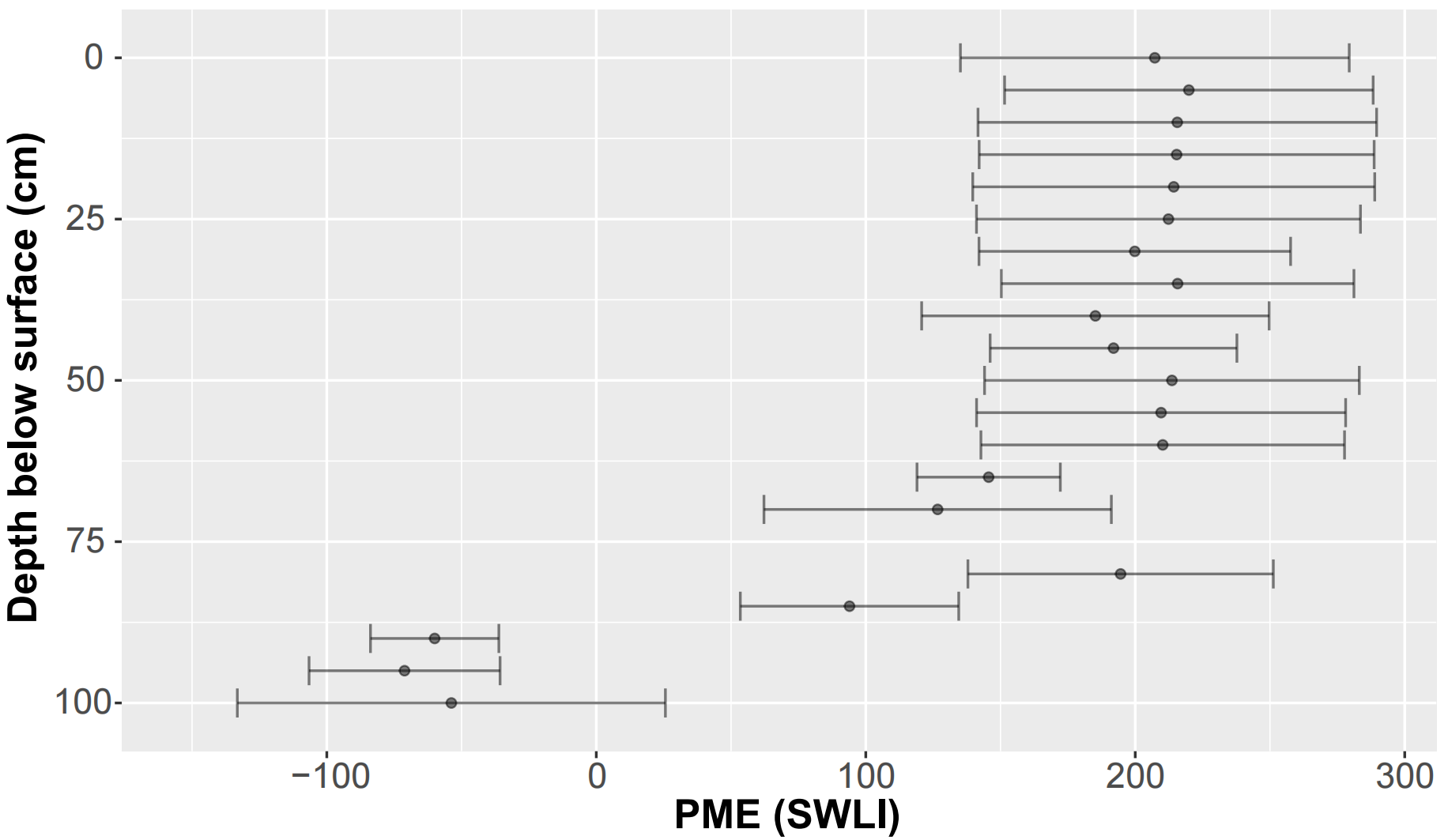

**Supplementary Fig. 5** Paleo-mangrove elevation (PME) reconstruction using morphology-based Bayesian transfer functions with *Ammonia tepida* grouped with *Ammonia* spp.. Predictions are derived from preserved foraminiferal assemblages in core MPSC01, with error bars indicating ±2σ sample-specific prediction uncertainties. Ammonia tepida was grouped with Ammonia spp. to further avoid misidentification risks. This conservative taxonomic approach led to exclusion of anomalous PME predictions in Unit III of MPSC01. Thus, this study retains morphological identification criteria remained consistent with established protocols for the study area (Supplementary Table 13)^43^.

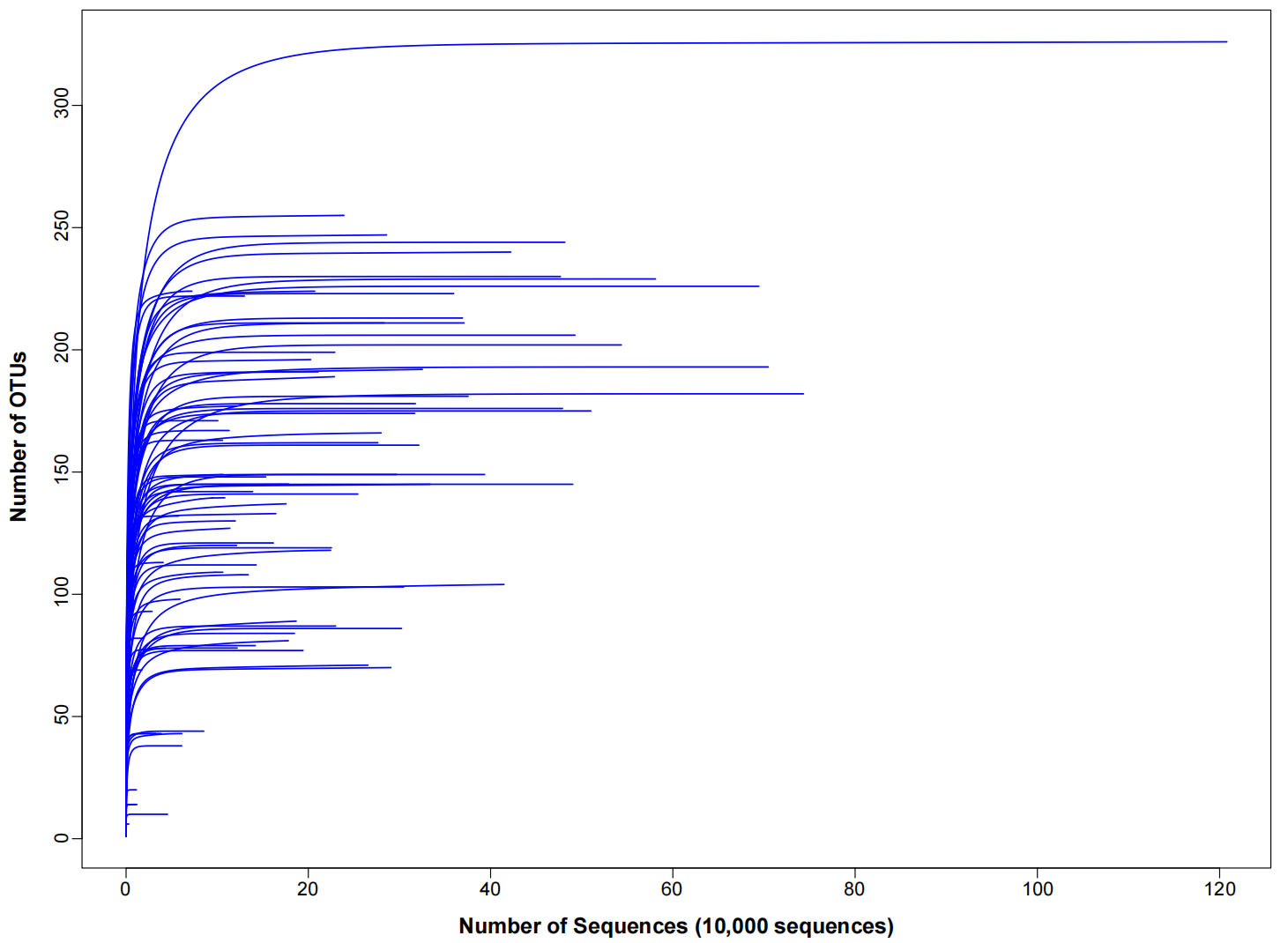

**Supplementary Fig. 6** Rarefaction of read counts against numbers of OTUs. Each curve represents a modern or core sample of the eDNA dataset.

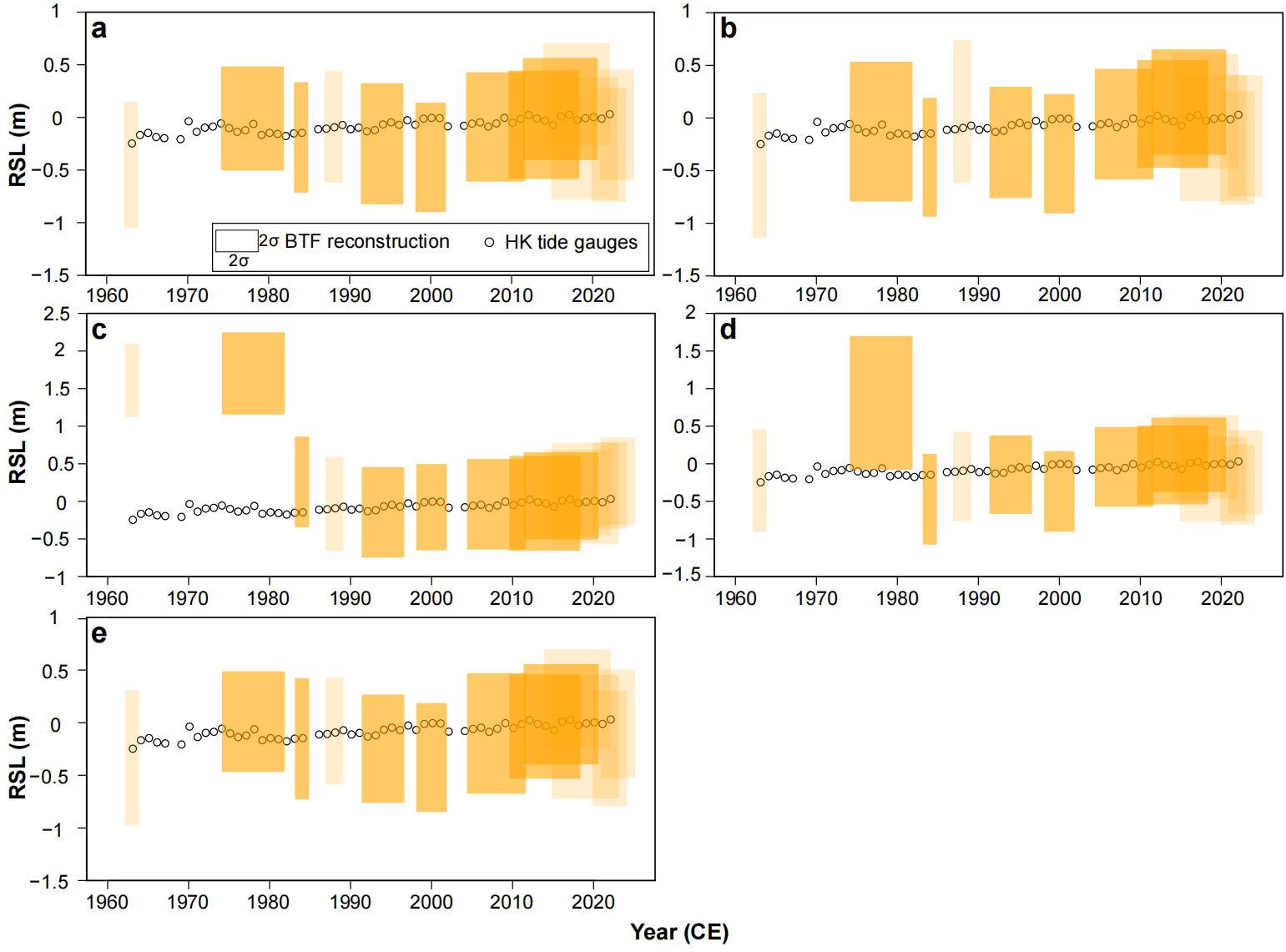

**Supplementary Fig. 7** RSL reconstructions and tide gauges records validation of core samples above 60 cm depth below surface using different combinations foraminiferal eDNA modern training sets for sensitivity tests. Boxes represent ±2σ vertical uncertainties and ±2σ age uncertainties for the reconstructions. RSL reconstructed from core samples with poor modern analogue were shown with transparent colors. Modern training set with all samples incorporated (a); Modern training set with only samples from mangrove environment (b); Modern training set with only samples from mudflat environment (c); Modern training set with only samples from MP_A and MP_B transects (d); Modern training set with samples from subtidal environment excluded are shown (e). Detailed performances of the tested datasets are provided in Supplementary Table 12.

In order to justify our choice of modern samples for eDNA-BTF modern training sets, we conducted a series of sensitivity tests on different sets of modern training sets. Our result suggests that the original eDNA modern training set with all modern samples included showed the highest performance, with the lowest average 1 σ uncertainty (18.16 SWLI units), a lowest root mean squared error of prediction (RMSEP; 24.5) and the strongest relationship between predicted and observed elevation (R^2^=0.95) in 10-fold cross-validation. Nevertheless, the mean squared error (MSE) of the validation against tide gauge records (0.032 m²) is slightly higher than that of the modern training set without subtidal samples (0.03 m²). This difference is likely due to the inclusion of eDNA samples from subtidal environments, which may introduce greater uncertainty when predicting the top section of the core—an interval that, according to LDA predictions, was deposited in a mangrove setting. Overall, to balance accuracy and interpretability, we retained our original eDNA modern training set.

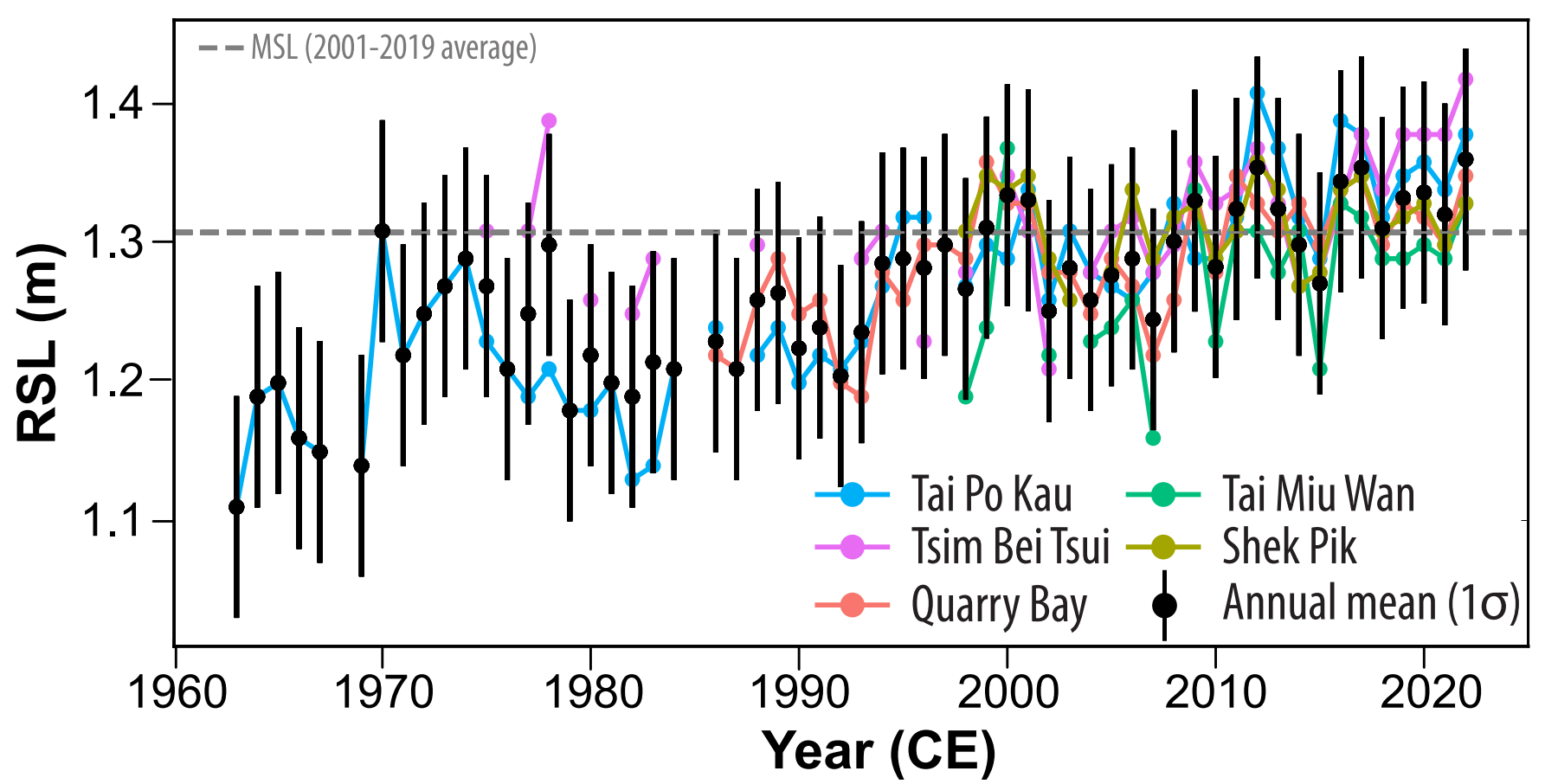

**Supplementary Fig. 8** Mean sea level changes (MSL ±1σ) relative to the 5-year (2017-2022) period average of all active tide gauge stations in Hong Kong. (Modified from Yu et al., 2025)^43^.
